# Ten-Fold Expansion MALDI Mass Spectrometry Imaging of Tissues and Cells at 500 nm Resolution

**DOI:** 10.1101/2024.10.20.619316

**Authors:** Chengyi Xie, Jianing Wang, Xin Diao, Lei Guo, Thomas Ka-Yam Lam, Ruxin Li, Yanyan Chen, Yue Zhang, Xiaoxiao Wang, Jiacheng Fang, Zongwei Cai

**Author notes:** Corresponding Authors: Dr. Jianing Wang; Prof. Zongwei Cai.

## Abstract

Achieving high spatial resolution in matrix-assisted laser desorption/ionization mass spectrometry imaging (MALDI-MSI) is crucial for detailed molecular mapping in biological tissues and cells. However, conventional MALDI-MSI platforms are typically limited to spatial resolutions of 5-20 μm, restricting their ability to visualize fine subcellular structures. Here, we present a novel ten-fold expansion MALDI-MSI (10X ExMSI) method that attains 500 nm spatial resolution without the need for specialized optics or customized instrumentation. By physically expanding biological samples using a swellable hydrogel matrix, our method maintains high retention and broadens the detection range of lipids, ensuring comprehensive lipid profiling. We demonstrated the effectiveness of 10X ExMSI by visualizing intricate subcellular structures within mouse brain tissues, including individual nuclei, astrocytes, Purkinje cells, and, notably, dendritic arborizations-features previously unresolvable using mass spectrometry imaging. Applying 10X ExMSI to single-cell analysis of cultured cells revealed detailed spatial distributions of lipids at the subcellular to organelle level. Notably, the method is fully compatible with standard commercial MALDI-MSI platforms, enabling widespread adoption without significant additional investment. The 10X ExMSI offers a transformative tool for high-resolution molecular imaging, opening new avenues for molecular analysis in diverse biological and biomedical fields, including neuroscience, pathology, and cellular biology.

## Introduction

Understanding the spatial distribution of biomolecules at the molecular level is fundamental to comprehending complex biological systems. High-resolution imaging techniques are essential for visualizing cellular components and elucidating biological processes within their precise spatial context. Optical microscopy, particularly super-resolution microscopy methods such as single-molecule localization microscopy, has significantly advanced our ability to observe structures beyond the diffraction limit, enabling visualization of organelles and sub-organellar features^1^. However, these optical imaging techniques primarily target proteins with antibodies that offer high specificity. While proteins can be precisely localized using antibodies, it inherently limits the scope of molecular imaging to those targets that antibodies can specifically recognize. Unlike proteins, imaging metabolites poses a significant challenge for optical microscopy due to the lack of suitable probes that offer the necessary specificity at the molecular level. Furthermore, the reliance on fluorescent dyes introduces additional limitations in thatthe spectral overlap significantly restricts the multiplexing capability and throughput of molecular imaging in a single experiment.

Matrix-assisted laser desorption–ionization mass spectrometry imaging (MALDI-MSI) provides label-free visualization of a wide range of biomolecules, including lipids, metabolites, and proteins, directly from tissue sections^2–4^. Unlike optical methods, MALDI-MSI does not rely on specific labels or dyes, thereby avoiding issues related to spectral overlap and limited specificity for small molecules. Despite its advantages, conventional commercial MALDI-MSI instruments are typically limited to spatial resolutions of 5–20 μm, which is insufficient for resolving tiny structures and investigating molecular compositions at the subcellular scale. To push forward the limitation, dedicated optical setups have been used to improve the resolution towards the low micrometer scale^5,6^. One way to improve the spatial resolution is adopting a transmission mode geometry, allowing the laser to be incident from the back of the slide, and to be focused on the sample surface^7–9^. This design achieved spatial resolution as high as 1.2 μm or even submicron level by oversampling^5^. Another way is creating a hollow lens to reduce the distance from lens to the sample surface, thereby decreasing the laser spot size and achieving resolutions around 1.4 μm^6,8^. However, these developed methods require sophisticated optics and customized instruments, thus limiting the broad applications of these methods. Additionally, these advanced optical setups have strict technical requirements for matrix deposition, requiring the use of sublimation methods. However, sublimation often leads to the decrease in sensitivity. Although techniques like secondary ion mass spectrometry (SIMS) and laser desorption/ionization (LDI) imaging can achieve spatial resolutions in hundreds of nanometers, they rely on hard ionization methods that result in extensive fragmentation of biomolecules. The fragmentation compromises the integrity of endogenous molecules, resulting in sparse and less informative biological data. Consequently, effective high-resolution biological mass spectrometry imaging relies on soft ionization techniques such as MALDI, which preserve molecular structures and provide abundant biological information. Enhancing the spatial resolution of soft ionization techniques, such as MALDI-MSI, is crucial for advancing molecular MS imaging capabilities.

Expansion microscopy (ExM) is an innovative solution that utilizes chemical methods to enhance spatial resolution without specialized microscopy to achieve super-resolution. ExM physically enlarges biological samples by embedding them in a swellable hydrogel, effectively increasing the spatial resolution of microscopy by the expansion factor of samples. This chemical method allows for nanoscale imaging using conventional microscopes. Since its introduction by Boyden and colleagues, ExM has been adapted and refined for various biological samples and imaging techniques.^10–14^. Building upon the principles of ExM, researchers developed expansion mass spectrometry imaging (ExMSI) methods for lipid imaging^15–18^, achieving a spatial resolution improvement of around 5-fold and attaining a lateral spatial resolution at 1 μm level^16^. Compared to optical microscopy techniques, which can achieve resolutions below 500 nm, existing MALDI-MSI methods—whether based on expansion or specialized instrument designs—have only reached spatial resolution limits around 1 μm. This significant gap highlights the need for further progress to bridge the resolution gap and attain nanoscale resolution in MALDI-MSI.

In this study, we introduce a novel ten-fold expansion MALDI-MSI (10X ExMSI) method that significantly enhances spatial resolution to approximately 500 nm. This method provides a spatial resolution comparable to that of optical microscopy while eliminating the need for specialized optics or customized instrumentation, making high-resolution MALDI-MSI more accessible and scalable. The 10X ExMSI method enables detailed molecular analyses at the nanoscale, allowing researchers to explore cellular heterogeneity and advance lipidomics research with unprecedented precision.

## Results and Discussion

### Development of 10x expansion-MSI workflow

To achieve high-resolution mass spectrometry imaging at the subcellular level, we developed a ten-fold expansion-MSI (10x ExMSI) workflow. The experimental details were described in the experimental section. Fig. 1a shows the experimental workflow of expansion-MSI. In brief, fixed tissue was cryosectioned into slides at 50 μm thickness and then the tissue slide was placed into a customed chamber consisting of two spacers. Then proteins are anchored to the gel matrix. Proteinase K (ProK) was used for tissue digestion at 60 ℃ in a humid environment to facilitate efficient expansion. After digestion, the gel was transferred to a large box to allow it to expand freely.

**Fig. 1:**
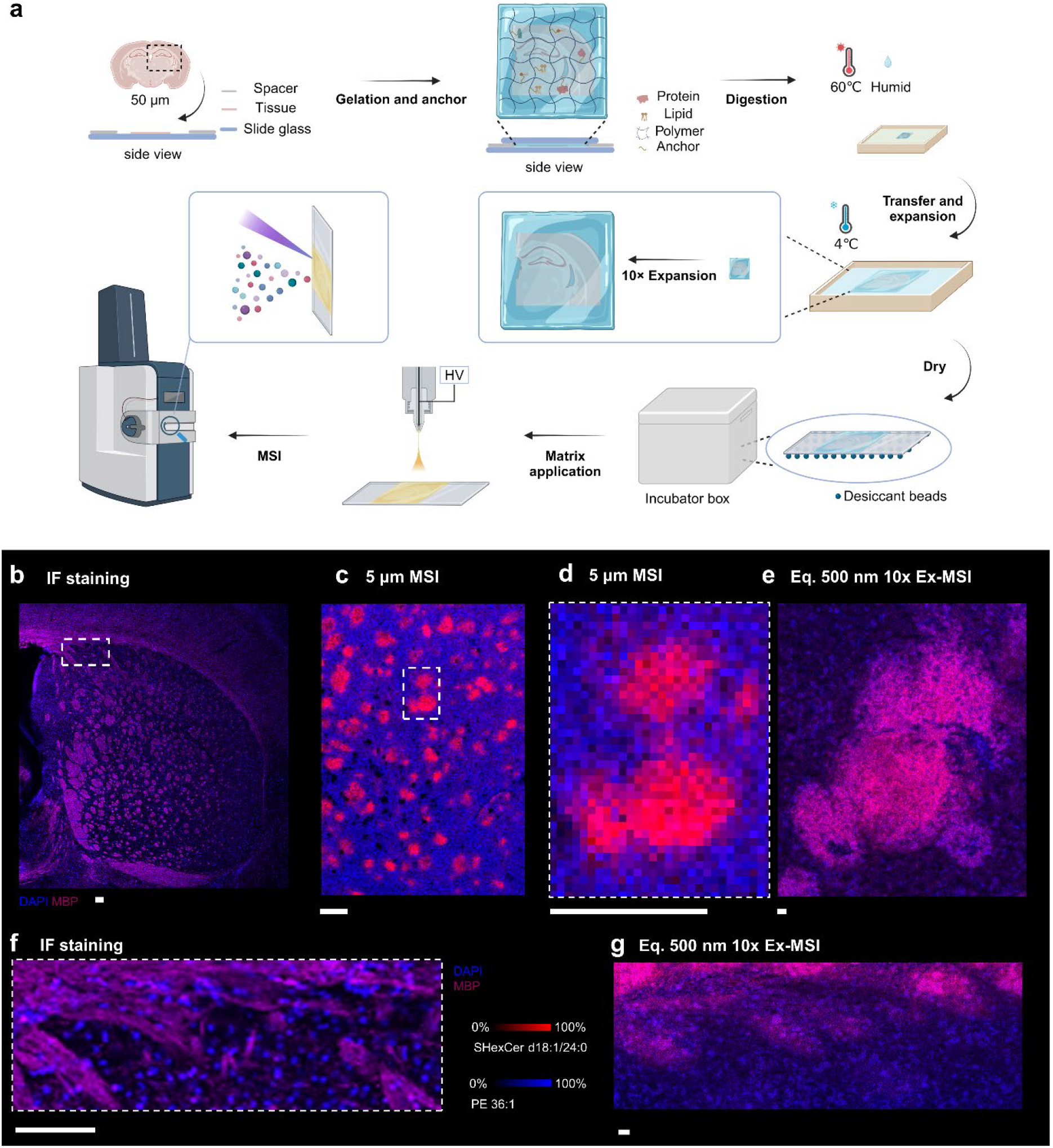
Schematic illustration of ten-fold expansion-MSI. a) step by step workflow of 10x ExMSI involved from sample preparation to data acquisition. b) immunofluorescence of myelin basic protein (MBP) in the fresh-frozen mouse brain at coronal plane. Nuclei were stained with 4’,6-diamidino-2-phenylindole (DAPI). c) ion images of m/z 888.62 (red) and m/z 744.56 (blue) for striatum region of fresh-frozen mouse brain under 5 μm MSI. d) Expended view of single caudate nucleus in the while boxed region in (c). e) ion images of m/z 888.62 (red) and m/z 744.56 (blue) for single caudate nucleus acquired by 10x ExMSI. An equivalent resolution of 500 nm was achieved under 5μm spatial resolution. f) Expended view of the while boxed region in (b). The region of the expanded view is similar to the region in (g). g) ion images of m/z 888.62 (red) and m/z 744.56 (blue) for part striatum region acquired by 10x ExMSI at an equivalent resolution of 500 nm. Scale bar represents 100 μm.

the orientation of the gel during expansion is crucial in our workflow. One critical step is that the gel should be flipped upside down with the tissue slice facing up or otherwise the signal of the sample will diminish during MSI acquisition since laser needs to penetrate the gel to reach the tissue plane. the gel was allowed to expand freely in deionized water at 4 ℃. The expanded gel then dried gently on the ITO slide placing in an incubator box filling with desiccant beads and gentle nitrogen gas flow. Fine matrix is then applied onto dry sample surface using a custom-built pneumatic-assisted electrospray deposition system to achieve high spatial resolution MSI.

We determined the expansion factor by measuring the dimensions of the embedded brain tissue before and after expansion (Supplementary Fig. 1). The bottom of the embedded brain tissue was measured at 6 mm, while the bottom of the same expanded sample was 61 mm after drying. The expansion factor (EF) could reach about 10.17-fold expansion using the developed 10X expansion MSI method. Therefore, as high as 500 nm equivalent spatial resolution can be achieved if 5 μm acquisition step is used for MALDI-MSI. The laser spot size is less than 5 μm to avoid oversampling (Supplementary Fig. 3).

The significant improvement in spatial resolution enables us to resolve subcellular structures that were previously unresolvable. The 500 nm equivalent spatial resolution provides the ability to differentiate nucleus in the striatum region in the overlay ion images of m/z 744.55 and m/z 888.62 (Fig. 1e and Fig. 1g), which were assigned as deprotonated PE 36:1 and deprotonated SHexCer d18:1/24:0 (herer d18:1 is assured as the major constituent of double-tailed sulfatide species), respectively. The distribution of nucleus can be aligned well with the DAPI staining results from conventional optical microscopy (Fig. 1f). In contrast, the same ion distributions obtained without expansion at 5 μm spatial resolution (Fig. 1d) fail to resolve the nuclear shapes.

The ten-fold ExMSI method thus effectively enhances the spatial resolution of MALDI-MSI beyond the limitations of conventional instrumentation, allowing for detailed molecular imaging at the subcellular level without the need for specialized optics or customized equipment.

### Fine-tuning Matrix Deposition for Nano-Scale Mass Spectrometry Imaging

Applying the matrix uniformly with appropriate crystal size is crucial for obtaining high-quality mass spectrometry imaging data, especially at high spatial resolution. α-Cyano-4- hydroxycinnamic acid (HCCA), N-(1-naphthyl) ethylenediamine dihydrochloride (NEDC), and 1,5-diaminonaphthalene (15DAN) showed good performance for high resolution metabolite analysis^20–22^, and therefore were adopted for 10X ExMSI. For a 5 µm step size, if the matrix crystal size is too large, it can not only cause molecule migration, which affects the effective resolution but also result in oversized laser crater. If the crater diameter exceeds 5 µm, oversampling occurs. To address this, we control the matrix crystal size to within 1.5 µm (Supplementary Fig. 2) to ensure uncompromised sampling. From the crater images, we can see that the laser-induced crater size is approximately 3.5 µm (Supplementary Fig. 3), leaving some margin beyond the requirements. In this scenario, we prioritize using a droplet-based matrix deposition system assisted by pneumatic electrospray which offers better sensitivity than sublimation-based methods. Our custom-built system meets the requirement of matrix application, and the detail was described in the experimental section.

One of the advantages of the ExMSI approach is that it places less stringent demands on matrix deposition conditions. The fine crystal size of matrix meets the MSI acquisition of timsTOF fleX with microGRID system at the highest spatial resolution of 5 μm. After the laser ablation, the laser ablated crater was measured less than 5 μm, indicating no oversampling was happened. In contrast, current imaging methods that utilize modified optical designes to achieve 1 µm resolution can only use sublimation with a limited selection of matrices, restricting measuring sensitivity^5^. Our developed method leaves significant room for further improvements in resolution in future work.

### High Coverage of Lipid Pathways with Enhanced Species Detection Following Ten-Fold Expansion

Mouse brain was used as an example model to demonstrate the effectiveness of 10X expansion MSI. Samples with and without expansion were both prepared for comparisons. Abundant ion signals are observed in the representative mass spectrum (Supplementary Fig. 4a) acquired in the expanded sample, which is consistent with the mass spectrum (Supplementary Fig. 4b) obtained in the fresh-frozen sample, indicating that the tissue expansion work flow did not result in significant lipid loss. In the positive ion mode, different distribution patterns are exhibited in the mass spectrum compared to the conventional sample (Supplementary Fig. 5). The dominant peaks changed from m/z 772.52 and 798.54 to m/z 756.55 and 782.57 after expansion. According to specie assignment results, both m/z 772.52 and 756.55 were assigned as PC 32:0, but were detected as [M + K]^+^ and [M + Na]^+^ ion forms, respectively. Similarly, m/z 798.54 and 782.57 were assigned to PC 34:0 as different alkali metal adducts as well. The expansion pretreatment leads to the dominant of sodium ion adducts detected for lipids in the positive ion mode, and therefore explaining the variation of distribution patterns in the mass spectra.

In total, 136 and 84 ion species were assigned in the positive and negative ion modes (Supplementary Table 1 and 2), respectively, within 5 ppm mass error using databases from LIPID MAPS. Among the assigned species, SHexCer, PE, an PA count in the top three in the negative ion mode (Supplementary Fig. 6), while PC, PS, and PI are assigned the most in the positive ion mode. We further compared the assigned species with fresh-frozen sections (Supplementary Table 3 and 4), which typically used for conventional MALDI imaging, to investigate specie lost after expansion. As shown in Supplementary Fig. 7, major loss is observed for PC and PE species in the positive and negative ion modes, respectively. However, the detection for other lipid species is beneficial from the expansion process, especially in the positive ion mode. The detection numbers of PS, PI, PG, and SHexCer are greatly increased compared to normal samples. The results demonstrate high lipid retention efficiency for 10X expansion MSI.

### Mouse Brain Expansion MSI

The fine structure of mouse hippocampus is closely related to its critical biological function, which highly demands high spatial differentiation capability to resolve the detailed structure. In this study, 20 μm MSI was performed for expanded mouse hippocampus and normal sample (Fig. 2a and 2d) to investigate the effective resolution increase using 10X ExMSI method. Overlayed ion images of m/z 834.53, m/z 885.55, and m/z 888.62 were exhibited to visualize the ion distribution at different spatial resolution. These species were assigned as deprotonated PS 40:6, PI 38:4, and SHexCer d18:1/24:0, respectively. In general, the major areas (alveus (alv), Cornu Ammonis (CA), and dentate gyrus (DG)) of the tissue are well preserved after expansion as compared to the fresh-frozen sample visualized by conventional MSI and confocal microscopy (Fig. 2a, 2b and 2c). The physical expansion improves the ability of MSI to resolve fine structure details in mouse brain tissues. While the overall morphology of the hippocampus can only be vaguely distinguished for the normal sample under 20 μm spatial resolution (Fig. 2a), more clear and sharp edge can be observed between different anatomical regions for the expanded mouse brain under the same laser spot size (Fig. 2d). The normal sample was additionally measured at 5 μm spatial resolution (Fig. 2b) to further prove the effectiveness of expansion MSI. While clearer ion images were obtained at 5 μm spatial resolution compared to 20 μm (Fig. 2a and 2b), the upper limitation spatial resolution of the instrument (i.e., 5 μm) still couldn’t resolve individual cells as shown in IF images in Fig. 2c. 10X ExMSI effectively improve the efficient spatial resolution of MSI to approach optical imaging. Comparing ion images in the expanded view of regions in Fig. 2e, it is apparent that the equivalent spatial resolution was increasing monotonously from 20 μm to 5 μm MSI and further to 20 μm ExMSI. The improvement provides the ability to discern tiny molecular difference at subcellular level. Specifically, the 20 μm ExMSI is readily resolve pyramidal cells in the pyramidal layer of field CA3 as well as cerebral cortex as shown in the expanded view (5) and (6) (Fig. 2e), while near harmonized ion distribution is observed in the corresponding area for the normal sample at 5 μm (expanded view (3) and (4)). The distribution of pyramidal neurons can be aligned to the staining results in view (7) and (8). Although single cell MSI for cultured cells was reported using commercial instrument, the differentiation of individual cells on tissue samples is more difficult due to the heterogeneity and overlapping of cells with different size^23^. In addition, the ion distribution shown as small-scale “strip” in view (6) acquired by ExMSI is consistent with the distribution of nucleus in white matter visualized by 4’,6-diamidino-2-phenylindole (DAPI) staining in view (8).

**Fig. 2:**
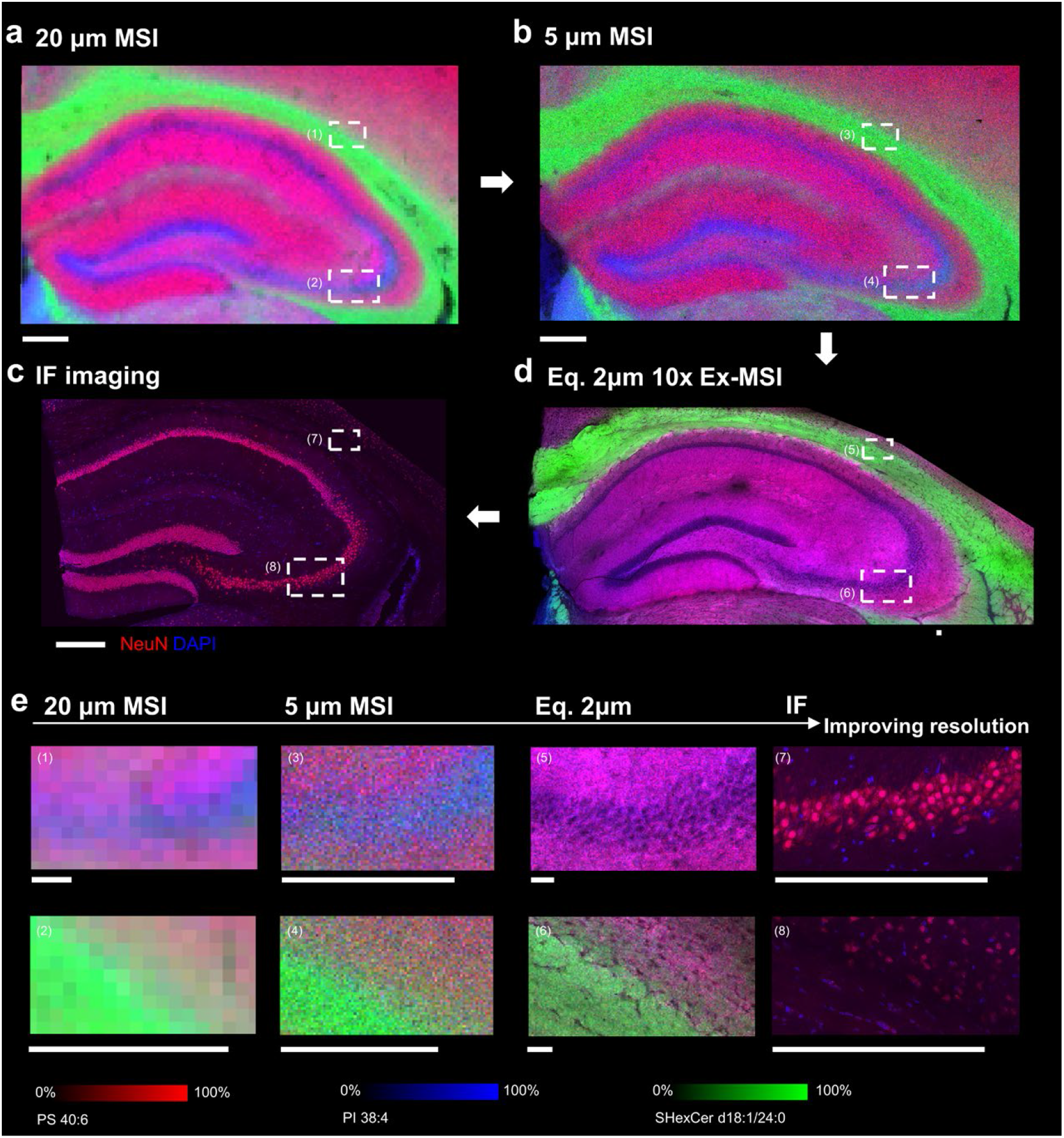
10X ExMSI of mouse brain hippocampus. overlay ion images of m/z 834.53 (red), m/z 885.55 (blue), and m/z 888.62 (green) acquired by a) 20 μm MSI, b) 5 μm MSI, and d) 20 μm 10X ExMSI. An equivalent spatial resolution of 2 μm was achieved for 10X EXMSI using 20 μm laser spot size. c) immunofluorescence of neuronal nuclei (NeuN) in the mouse hippocampus. Nuclei were stained with 4’,6-diamidino-2-phenylindole (DAPI) as well. e) expended view of boxed regiones in (a-d). Scar bar represents 300 μm.

To further prove the effectiveness of the expansion MSI method, mouse cerebellum was tested as well and the corresponding expansion MSI images were shown in Supplementary Fig. 8c corresponding to m/z 718.54, and m/z 888.62, which are assigned to PE 34:0 and SHexCer d18:1/24:0 as deprotonation ion form. The major areas of white matter layer, granule layer, and molecular layer are fully resolved in the overlay ion image. When compared to normal sample acquired at 5 μm spatial resolution (Supplementary Fig. 8b), similar to the results obtained from the mouse hippocampus sample, clearer ion images were obtained for expanded mouse cerebellum under a spatial resolution of 20 μm with a four-fold difference in laser spot size. In detailed, the granular neuron cells are neatly arranged in the region (4) acquired in optical image, but the 5 μm MSI is unable to differentiate the detailed ion distributional differences in the granular layer in region (2) which are shown as scatted pixels. The 20 μm 10X ExMSI provides better visualization for the fine structure in the granular layer, in which neurons cells can be partially discerned in the overlay ion image. The distribution of neurons cells can be aligned to the staining results using luxol fast blue (LFB) and cresyl violet (CV) in region (4). Since nearly 10X expansion factor was achieved, it is not surprising that more effective spatial differentiation capacity was achieved with only a 4-fold difference in laer spot size. Along with hippocampus and cerebellum, 20 μm ExMSI was performed for the olfactory areas of mouse brain (Supplementary Fig. 9). The overlay image of species [PE 34:0-H]^-^, [SHexCer d18:1/24:0-H]^-^, and [PI 34:0 - H]^-^ reveals sophisticated anatomic structure of the olfactory areas. A thin sheet composed of numerous spherical glomeruli in the outer layer of olfactory bulb is visualized, which is related to the odor map, while the 20 μm conventional MSI only resolved a thin sheet without any detailed structure information. Additionally, a densely packed and extensive distribution of granule cell layer is visualized in the ion species of [PE 34:0-H]^-^ and [SHexCer d18:1/24:0-H]^-^ by 10X ExMSI. The results of expansion MSI for hippocampus, cerebellum, and olfactory bulb confirm that the 10X ExMSI far exceeds the upper spatial resolution limitation of commercial instrument, enabling on-tissue profiling of single cells at subcellular level under a soft spatial resolution of 20 μm.

### High spatial resolution expansion MSI

10X ExMSI was readily resolved the detailed structure of mouse hippocampus both in the negative and positive ion modes (Fig. 2d and Supplementary Fig. 10) under a soft spatial resolution of 20 μm. To investigate the biological structural discrimination ability using higher spatial resolution, 10 μm MSI was performed for part region of mouse hippocampus and cerebellum after expansion. Fig. 3a shows overlay ion images of PS 40:6, SHexCer d18:1/24:0, PI 38:5, and m/z 844.41. The first three species were detected as [M-H]^-^ ion form at m/z 834.53, 888.62, and 883.53, respectively. m/z 844.41 was unassigned based on a mass tolerance of 5 ppm using database from LIPID MAPS. In addition to differentiation of individual neuron cells proven above, the combination of 10 μm laser spot and 10X expansion enables the positioning of astrocytes across the hippocampus section in the overlay ion image, which were appeared as strip or star shape (Fig. 3a). The distribution and shape of astrocytes visualized by ExMSI is consistent with the IF of the glial fibrillary acidic protein (GFAP) staining (Fig. 3b). Additionally, high spatial resolution ExMSI resolves subcellular structure for mouse cerebellum. The distributions of m/z 599.32, 718.54, and 888.63 are displayed in Fig. 3c and assigned to the [M – H]^-^ ion form of SHexCer d18:1/24:0, LPI 18:0, and PE 34:0. Our ExMSI data revealed that SHexCer d18:1/24:0 is dominated in the white matter, which is mainly made up of myelinated axons, and decrease rapidly in the granule layer. Notably, the dotted distribution of nucleus is revealed, where most of them are in the granule layer and a few in the white matter layer, in the ion image of LPI 18:0 employing 10 μm ExMSI. The nucleus mainly distributed inside granular neuron cells in the granular layer, which is according with IF results of DAPI and MBP. A complementary ion distribution to LPI 18:0 can be observed for the ion specie PE 34:0, whose distribution further extends into the molecular layer. The successful visualization of individual nucleus is owing to the high spatial resolution achieved by 10X ExMSI, leading to the ability to resolve subcellular details at molecular level.

**Fig. 3:**
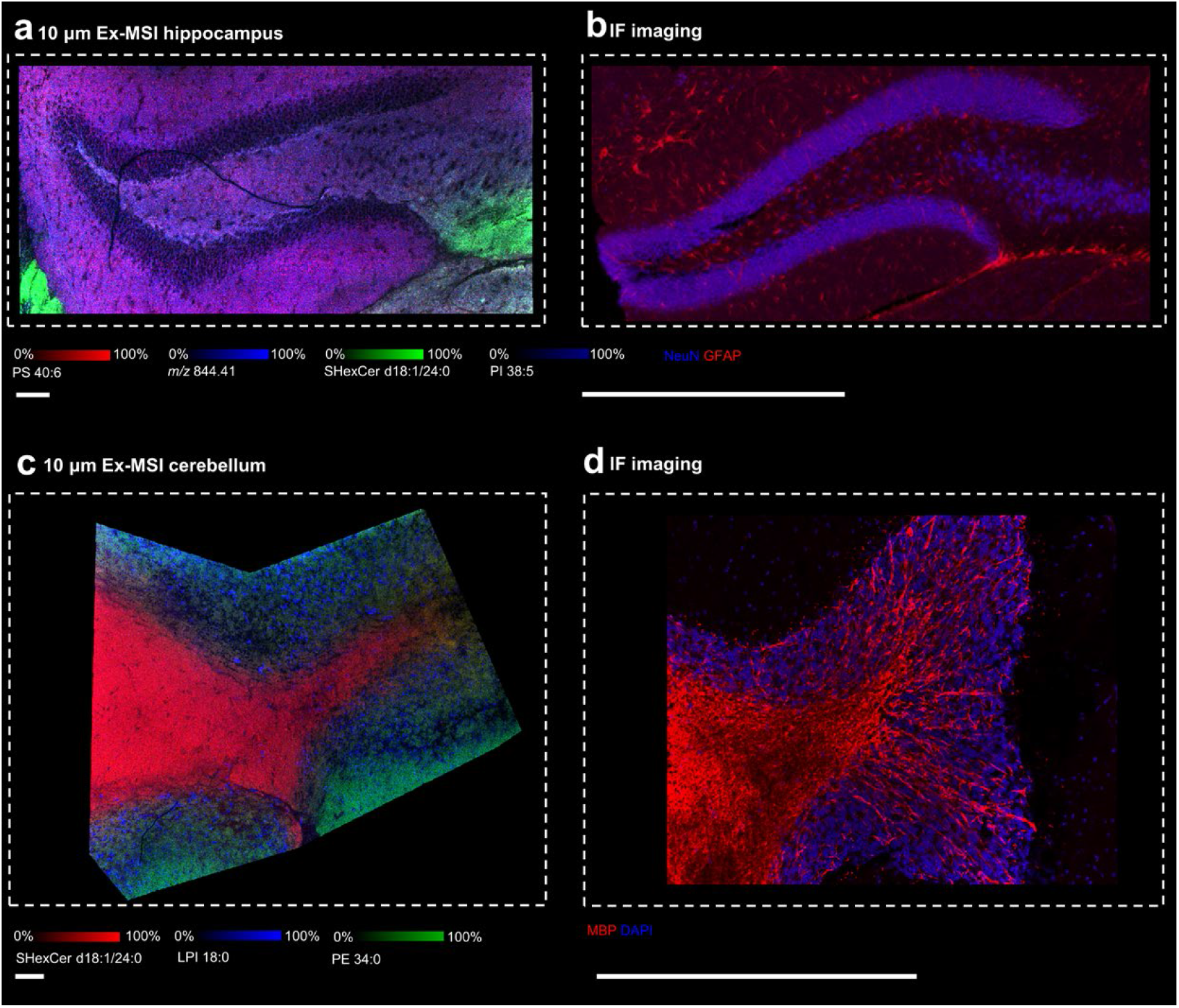
high resolution 10X ExMSI. a) overlay ion images of m/z 834.53 (red), m/z 844.41 (light blue), m/z 888.62 (green), and m/z 883.53 (dark blue) for mouse hippocampus acquired by 10 μm 10X ExMSI. An equivalent spatial resolution of 2 μm was achieved for 10X EXMSI using 20 μm laser spot size. b) immunofluorescence of neuronal nuclei (NeuN) and Glial fibrillary acidic protein (GFAP) in the mouse hippocampus. c) overlay ion images of m/z 888.62 (red), m/z 599.32 (blue), and m/z 718.54 (green) for mouse cerebellum acquired by 10 μm 10X ExMSI. d) immunofluorescence of MBP in the mouse cerebellum. Nuclei were stained with 4’,6-diamidino-2-phenylindole (DAPI) as well. Scar bar represents 500 μm.

To characterize the equivalent lateral resolution of ExMSI, we next performed experiments for mouse cerebellum with and without expansion in which laser spot size was varied from 100 μm to 5 μm (Fig. 4b and Fig. 4c). Adjacent parallel regions were selected to keep other experimental parameters unchanged. Fig. 4a displays the distributions of ion species at m/z 751.52, and 756.55 and assigned as [M+Na]^+^ adducts of PC 32:0, and PA 38:2. Major areas of cerebellums, including white matter layer, granular layer, Purkinje layer, and molecular layer, are readily differentiated using 20 μm positive ion mode 10X ExMSI. PC 32:0 is highly expressed in the granular layer and molecular layer, while PA 38:2 is enriched in the white matter which is mainly made up of nerve tracts. In particular, a series of aligned Purkinje cells are observed as red spots in the Purkinje layer visualized in the ion image of PA 38:2. The oval morphology of Purkinje cells is clearer when employing higher spot size (10 μm and 5 μm) in Fig. 4b. Notably, part of the dendrite system of Purkinje cells is visualized as red lines connected to cell bodies under high resolution ExMSI (Fig. 4d). To clearly identify the dendrite system from the brain, the quality of the region of interest (ROI) was enhanced using colocalized ions (Fig. 4f). Details of the enhancement process is described in the Methods section. The distribution of the dendrite system can correspond to the IF staining results (Fig. 4e). Considering the 100 nm to 1 μm in width of the complex dendrite system before expansion, it is possible to resolve them after tenfold expansion, strongly demonstrating submicron resolution was achieved using the developed 10X expansion MSI method. According to our knowledge, the visualization of dendrite system is firstly reported in the area of MS-based imaging. Although cell body of Purkinje cells could be visualized by employing transmission- mode geometry MALDI (t-MALDI) and laser-induced postionization (MALDI-2) under 1.1 μm spatial resolution^5^, the dendrite system is still undifferentiable. Additionally, the submicron spatial resolution previously reported visualized microfissures and cracks in tissue slides resulted from sample pretreatment ^5^, resulting in a decrease in the actual effective spatial resolution. In this study, the small-scale fissures haven’t been observed possibly due to the protection of hydrogel to reduce cracks and fissures of tissue slides.

**Fig. 4:**
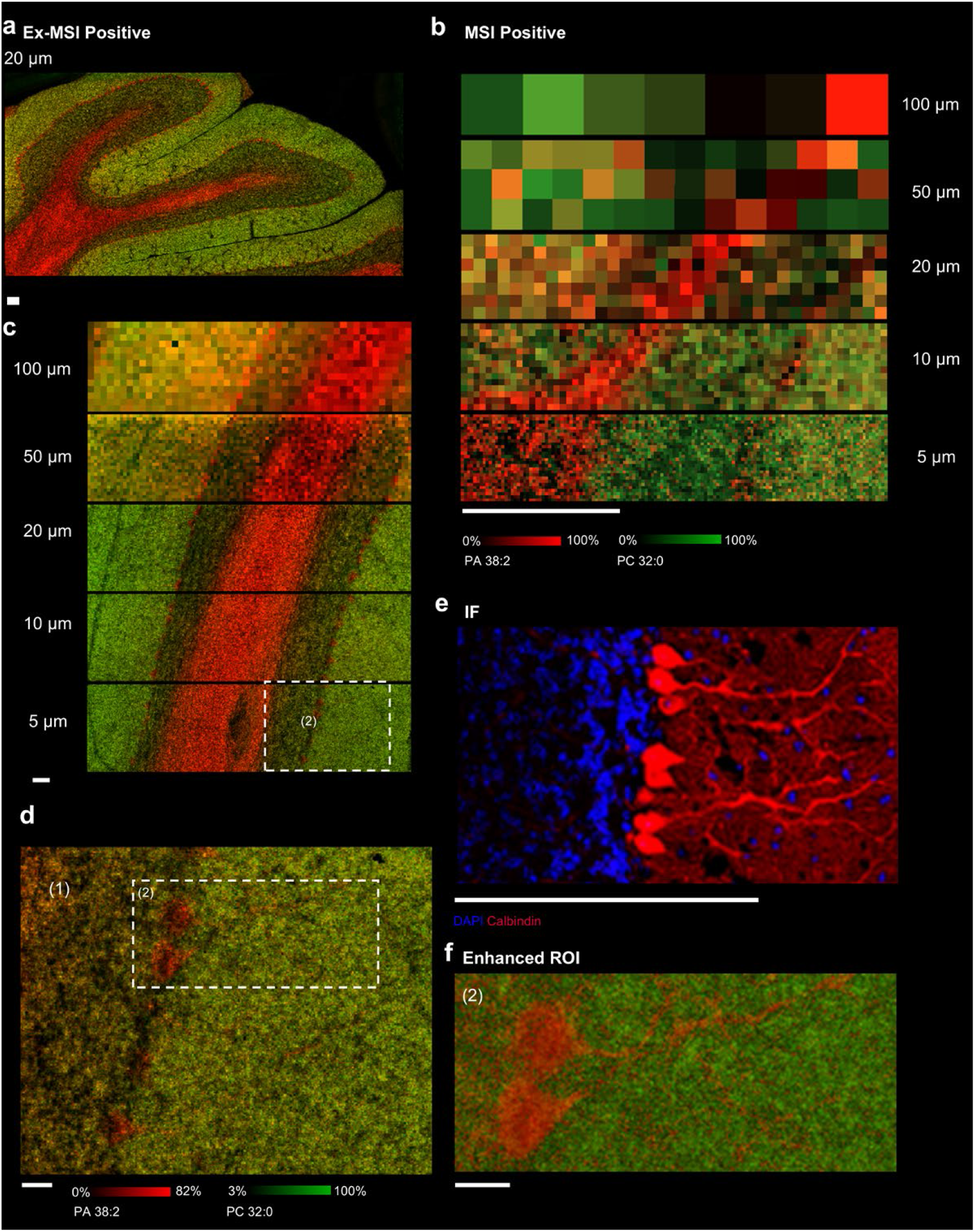
high resolution 10X ExMSI for mouse cerebellum. a) overlay ion images of m/z 751.52 (red) and m/z 756.55 (green) for mouse cerebellum acquired by 20 μm 10X ExMSI. b, c) overlay ion images of mouse cerebellum acquired by laser spot size was varied from 100 μm to 5 μm using (b) 10X ExMSI and (c) conventional MSI. d) expended view of boxed regiones in (b). e) immunofluorescence of calbindin in the mouse cerebellum. Nuclei were stained with 4’,6-diamidino-2-phenylindole (DAPI) as well. f) Enhanced region of interest (ROI) of the boxed region in (d) after data processing. The details were listed in the Methods. Scar bar represents 250 μm.

For images acquired by ExMSI, the decrease in spot size provides clearer visualization of the fine structure of the Purkinje layer. It is apparent that the greatly enhancement of equivalent lateral resolution for ExMSI when compared to unexpanded sample acquired at the same lateral resolution. With the increasing of sampling pixel size, although the distribution of dendrite system is becoming ambiguous leading to ‘scattered’ pixels across the molecular layer, a thin layer of the cell body of Purkinje cells is still discernible after expansion even under the spatial resolution of 100 μm (Fig. 4c). However, the 100 μm spatial resolution for conventional MSI only provide several pixels failing to form a distinguishable ion image (Fig. 4b). In addition, the cell bodies of Purkinje cells are difficult to be distinguished when the spot size exceeds 10 μm for unexpanded sample. Even when comparing to 5 μm MSI, 50 μm ExMSI could provide equivalent or better visualization of Purkinje layer, proving the effectiveness of ten-fold ExMSI achieved in this study.

### Single Cell ExMSI

To explore the potential of our developed 10X ExMSI method for the spatial lipid analysis of single cells, A549 cells were investigated in this study. Fig. 5a shows the representative mass spectrum acquired at 5 μm spatial resolution. Abundant ion signals could be observed in the spectrum and can be assigned to different kinds of lipid species (Supplementary Table 5 and 6). The ion distributions of assigned ion species of protonated SM d18:1/16:0, sodiated PC 34:1, and m/z 666.43 are exhibited in Fig. 5b expressed at distinct subcellular distribution. While SM d18:1/16:0 and PC 34:1 are basically co-located across cytoplasm and nucleus, the unknow species at m/z 666.43 has extended distribution across the plasma membrane. These two-ion distribution possibly located the organelle distribution of mitochondria and endoplasmic reticulum within cells by comparing the IF results (Fig. 5e). In the comparison of the ion images obtained from normal samples (Fig. 5c), the ion distribution is preserved after expansion, and the morphology of the cells is well displayed in that all structure of the cell body is clearly visualized after expansion using 5 μm laser spot size in the overlay ion image. In detail, the cavity shape of the nucleus can be visualized after expansion while the location of nucleus is unable to be resolved for unexpanded cells using the same laser spot size of 5 μm. As shown in the expanded view in Fig. 5d, the ion distribution is not precisely localized due to the limited spatial resolution as compared to ExMSI. The above results demonstrate the developed ExMSI method can be applied for the spatial lipid analysis of single cells, enabling the explanation of organelle structure, which is important in understanding cell-to-cell variability and the specific roles of lipids in different cell types or even in different states of the same cell type.

**Fig. 5:**
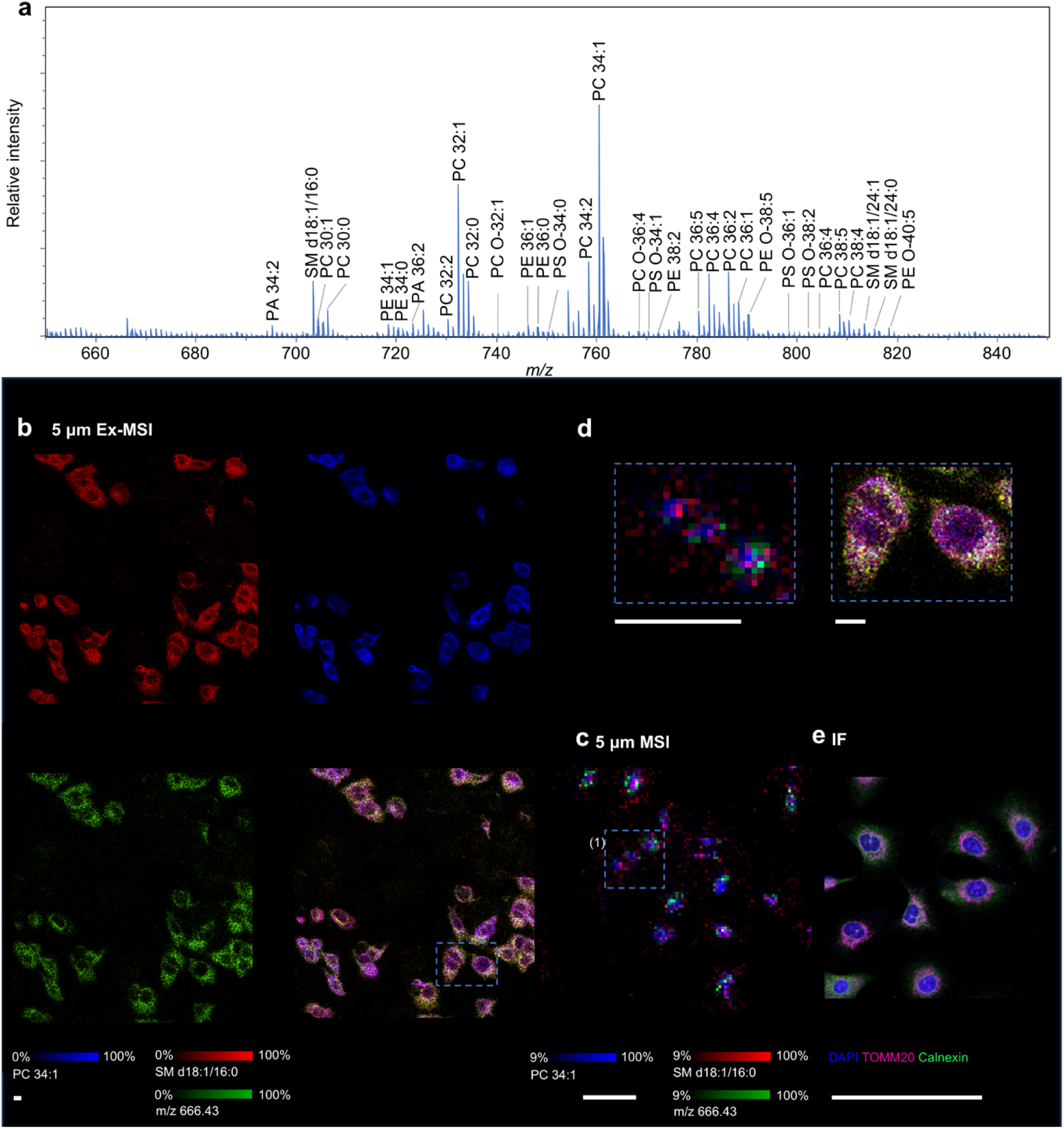
Single cells expansion mass spectrometry imaging. a) Averaged mass spectrum of ExMSI for A549 cells acquired at 5 μm pixel size in the positive ion mode. b) Ion images of m/z 703.57 (red), m/z 760.58 (blue), and m/z 666.43 (green) for A549 cells by ExMSI acquired at 5 μm pixel size in the positive ion mode. c) Ion images of m/z 788.54 (blue), m/z 833.53(green), and m/z 885.55 (red) for A549 cells by normal MSI acquired at 5 μm pixel size in the positive ion mode. d) expanded view of boxed regiones in (b) and (d). e) immunofluorescence of TOMM20 and Calnexin in the mouse cerebellum. Nuclei were stained with 4’,6-diamidino-2-phenylindole (DAPI) as well. In summary, the combination of ten-fold expansion and high resolution MSI (5 μm spatial resolution) enables us to visualize spatial lipidomics into a smaller scale using equivalent submicron spatial resolution.

### Conclusion

A ten-fold expansion mass spectrometry imaging (10x ExMSI) was developed for lipid mass spectrometry imaging using existing commercially available mass spectrometer without any modification. Under the confines of a 5 μm spot size, an equivalent spatial resolution of 500 nm was achieved by MALDI MSI using 10x ExMSI. The 10x ExMSI enables detailed imaging of biological samples’ structures. With this advanced method, individual cells can be distinctly identified in mouse brain tissue. Notably, the dendritic network of Purkinje cells, with widths spanning from 100 nm to 1 μm, was successfully imaged using 10x ExMSI. Furthermore, this approach allows for the visualization of subcellular structures within single cells. The developed 10x ExMSI method paves critical way for deciphering biological function of lipids at subcellular level.

## Methods

### Reagents and materials

| Reagent | Acronym | Vendor | Catalog Number |
| --- | --- | --- | --- |
| Sodium acrylate | SA | Santa Cruz Biotechnology | sc-236893D |
| Sodium acrylate | SA | AK Scientific | R624 |
| N,N-Methylenebisacrylamide | BIS | Sigma Aldrich | M7279 |
| N,N-dimethylacrylamide | DMAA | Sigma Aldrich | 274135 |
| Acrylamide | AA | Sigma Aldrich | A8887 |
| 4-hydroxy-TEMPO | 4HT | Sigma Aldrich | 176141 |
| N,N,N',N'Tetramethylethylenediamine | TEMED | Sigma Aldrich | T9281 |
| Methacrolein |  | Sigma Aldrich | 133035 |
| Phosphate Buffered Saline, 10x Solution, PH 7.4 | PBS | Thermo Fisher Scientific | 70011044 |
| Ammonium persulfate | APS | Sigma Aldrich | A3678 |
| Sodium chloride | NaCl | Sigma Aldrich | S6191 |
| Triton X-100 |  | Sigma Aldrich | T8787 |
| Proteinase K (PCR grade) | ProK | Thermo Fisher Scientific | EO0492 |
| Tris-HCl, 1 M, pH 8 |  | Thermo Fisher Scientific | 15568025 |
| 2,5-dihydroxybenzoic acid | DHB | Sigma Aldrich | 149357 |
| N-(1-naphthyl) ethylenediamine dihydrochloride | NEDC | Sigma Aldrich | 222488 |
| 1,5-diaminonaphthalene | 15-DAN | J&K Scientific | 209271 |
| 4% Paraformaldehyde Fix Solution |  | Beyotime Biotechnology | P0099 |
| sucrose |  | Sigma Aldrich | S112228 |
| Superfrost Plus Adhesion Slide |  | Epredia | J1800AMNZ |
| Indium tin oxide-coated glass slides (25mm x 75 mm) | ITO | Delta Technologies | CG-90IN-S107 |
| Indium tin oxide-coated glass slides (50mm x 75 mm) | ITO | Delta Technologies | CG-90IN-S207 |
| Methanol (HPLC-grade) | MeOH | VWR | BDH20864 |
| 4',6-diamidino-2-phenylindole | DAPI | Thermo Fisher Scientific | D1306 |

### Animal

C57/BL mice were purchased from Laboratory Animal Service Centre at the Chinese University of Hong Kong. They were housed in standard conditions with 12-hr light/dark and fed with sterilized water and standard laboratory feed. All animal experiments were approved by the Hong Kong Baptist University Committee on the Use of Human and Animal Subjects in Teaching and Research.

### Preparation of tissue slides

The mice were sacrificed, and organs were dissected and postfixed using 4% PFA with 0.1% glutaraldehyde in 1x PBS overnight at 4 °C. They were washed using PBS overnight at 4 °C and then cryoprotected by incubating in 15% and 30% (w/v) sucrose in 1x PBS at 4 °C until they sank. Prepared tissue was stored at -80 °C before section. 50 μm sections were prepared for expansion and immunofluorescence using CryoStar Nx70 cryostat (Thermal Fisher Scientific, Walldorf, Germany) at a chamber temperature of -20 °C. 10 μm sections were prepared for nonexpansion slides at the same section conditions.

### Gelation

A monomer solution was freshly prepared consisted of 34% SA (w/v), 10% AA (w/v), 4% DMAA (v/v), 1% NaCl (w/v), 0.01% BIS (w/v), and 1× PBS. The prepared mixture was stored at 4 ℃ before use. Before gelation, tissue slices were placed into custom gelling chamber consisting of two spacers cut from no. 1.5 cover glass adhered to a microscope slide. Excess PBS around the tissue was absorbed and sections were allowed to air dry on the slide. The tissue slices were transferred to the gelation chamber. Immediately before gelation, the chemicals 4HT, TEMED, APS and methacrolein were added to the gel monomer solution to a final concentration of 0.25% (w/v) APS, 0.001% 4HT (w/v), 0.04% TEMED (v/v), and 0.1% (v/v) methacrolein, adding TEMED and APS last to prevent premature gelation. Slides were incubated in gelling solution for 30 min at 4 °C to allow the monomer solution to diffuse into the tissue. A glass microscope slide wrapped with parafilm was placed over the gelling chamber and the samples were incubated overnight in a humidified container at 37 °C to complete gelation.

### Expansion

Excess gel around the tissue slice was trimmed off into an asymmetric shape. The gelation chamber was put into tailor-made ITO slide box filled with 6 units/mL proteinase K in digesting solution composed of 50 mM Tris-Cl (pH 8), 25 mM EDTA (pH 8), 0.5% Triton-X 100, 0.8 M NaCl, and Digestion was carried out at 60 °C for 1.5 hr in a humid chamber. After digestion, the gelation chamber was taken out and poly-lysine coated ITO slide was placed inside the box with the conductive side facing up. The gel was detached from the gelation chamber and flipped upside down with the tissue slice facing up using a fine brush. The gel was rinsed by 4 °C PBS for 15 mins three times on ice. The gel was then expanded in 4 °C deionized water for 20 min four times on ice, until there was no further expansion. The gel was rinsed by 4 °C deionized water three times quickly on ice. After rinsing, liquid was removed as much as possible and the ITO slide box was put into a drying chamber filled with silica gel desiccant beads under gentle nitrogen gas flow until fully dry.

### Immunofluorescence (IF)

Prepared tissue sections were washed three times using 1X PBS. The sections were permeabilized by 0.5% Triton X-100 in 1X PBS for 20 min. Tissue sections were washed twice using 1X tris-buffered saline (TBS). The sections were incubated in the mixture of 5% goat serum and 1% bovine serum albumin (BSA) for 2h and then incubated with primary antibodies overnight at 4 °C. Tissue sections were washed three times using 1X TBS and incubated for 3 min with DAPI. Images were captured by an DMi8 confocal microscope system (Leica, German). The following primary antibodies were used: Alexa Fluor 488-conjugated anti- Myelin Basic Protein (MBP) Antibody (850908, Biolegend), Alexa Fluor 568-conjugated Anti- neuronal nuclei (NeuN) antibody (EPR12763, abcam), Alexa Fluor 488-conjugated anti- Glial fibrillary acidic protein (GFAP) Antibody (644704, Biolegend), and Alexa Fluor 594- conjugated Anti-Calbindin Antibody (88831, Cell Signaling Technology).

### Matrix application

HCCA (10 mg/mL in 30% H2O and 70% MeOH), NEDC (5 mg/mL in 30% H2O and 70% MeOH), and 1,5-DAN (5 mg/mL in 30% H2O and 70% MeOH) were freshly prepared before matrix application. Matrix was applied using a custom-built matrix deposition device with the settings as the followings: N2 sheath gas (99.995 %) 70 psi, temperature 60 °C, matrix flow rate 10 µL/min, capillary voltage 5000 V, XY-stage speed 10.98 cm/min, line-to-line distance of 1 mm, spray layers 10 cycles.

### MALDI analysis

MSI experiments were performed on a timsTOF fleX MALDI-2 (Bruker Daltonics, Bremen, Germany) equipped with microGRID. Lateral resolution was set to 5-50 μm with 60-90% laser power in single mode at 10000 Hz and 20-200 shots per pixel. Mass range was set to m/z 100- 2000. Mass calibration was performed before the start of each MSI experiment using the ESI- L Low Concentration Tuning Mix (Agilent Technologies, CA, USA). Acquired MSI data was visualized using SCILS Lab MVS (Bruker Daltonics, Bremen, Germany).

### Ion specie assignment

Specie assignment of identified ion peaks was based on standard library search using LIPID MAPS® Structure Database (LMSD) from LIPID MAP with a mass tolerance of 5 ppm. Ion adduct forms of [M + H]^+^, [M + Na]^+^, [M + K]^+^, and [M + 2Na - H]^+^, and [M - H]^-^, and [M + Cl]^-^were considered for the positive and negative ion modes, respectively. The consideration of [M + Cl]^-^ form in the negative ion form is due to the exist of HCl in the NEDC and this ion from was reported in previous research^20^. To deduce false positive rate, only fatty acyl chains with even length was considered. Artificial screening was conducted to filter non-mammalian and unreported species.

### Enhancing ROI Quality

To clearly identify the detailed structure from the brain, data processing is conducted in the following three steps:

Firstly, the proposed DeepION model^24^ is employed to identify four colocalized ions for the query ions *m/z* 751.526 and *m/z* 756.552, as displayed in Supplementary Fig. 11A.Secondly, these ten ions, including *m/z* 751.526, *m/z* 810.599, *m/z* 811.603, *m/z* 856.580, *m/z* 797.507, *m/z* 756.552, *m/z* 782.566, *m/z* 783.570, *m/z* 757.555 and *m/z* 753.588 are combined, and the dimensionality reduction algorithm, Uniform Manifold Approximation and Projection (UMAP), is applied to obtain 3-dimensional embedding data, as shown in Supplementary Fig. 11B.Thirdly, the embedding data (Supplementary Fig. 11B) is then fused with the original data (Supplementary Fig. 11C) by weighted summation to obtain the enhanced dataset, as shown in Supplementary Fig. 11D.

## Conflicts of Interest

There are no conflicts to declare.

### Acknowledgments

This work was supported by General Research Fund (12302122) of the Research Grants Council, Hong Kong Special Administrative Region, SKLEBA Research Grant (SKLP_2021_P04), and a Start-up Grant from Hong Kong Baptist University.

A US provisional patent application has been filed covering the technology/methodology described in this manuscript.

