## Supplemental Information for "Ten-Fold Expansion MALDI Mass Spectrometry Imaging of Tissues and Cells at 500 nm Resolution"

[*] Corresponding Authors:

Dr. Jianing Wang

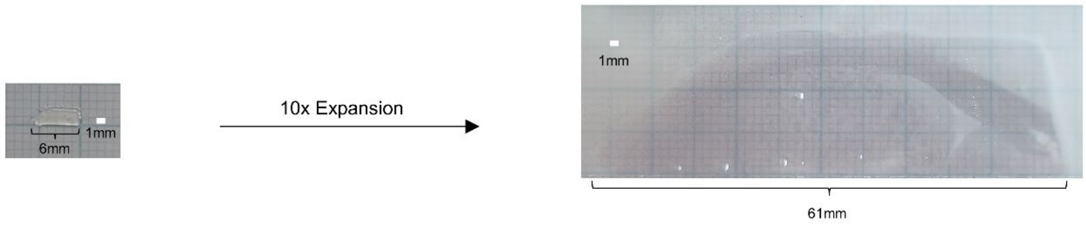


**Supplementary Fig. 1**: Measurement of the bottom of the section before and after expansion.

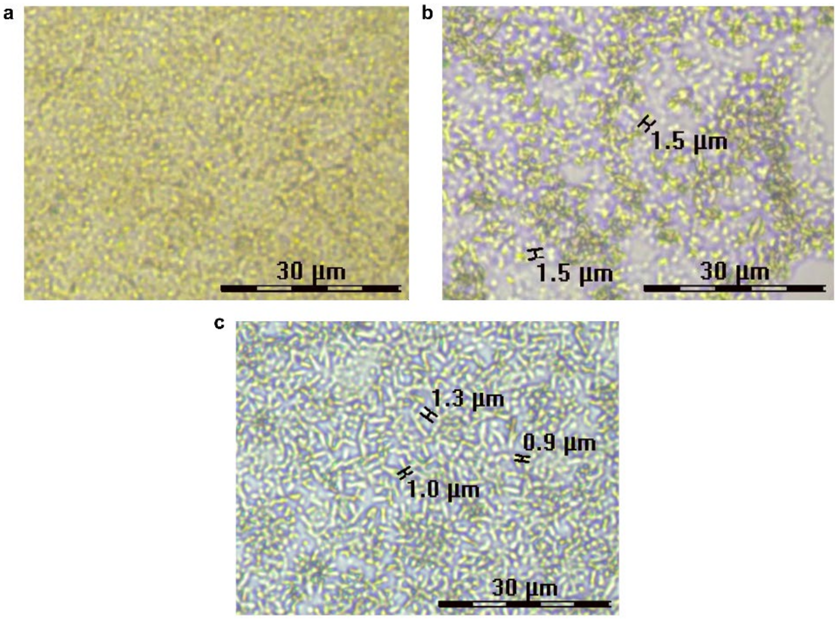


**Supplementary Fig. 2**: Measurement of crystalline size for matrix after the application of a) HCCA, b) NEDC, and c) 15-DAN.

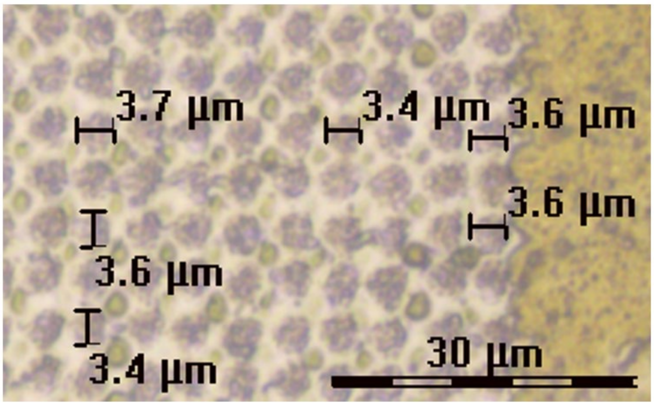


**Supplementary Fig. 3**: Measurement of ablated craters after the ablation under 5 μm spatial resolution.

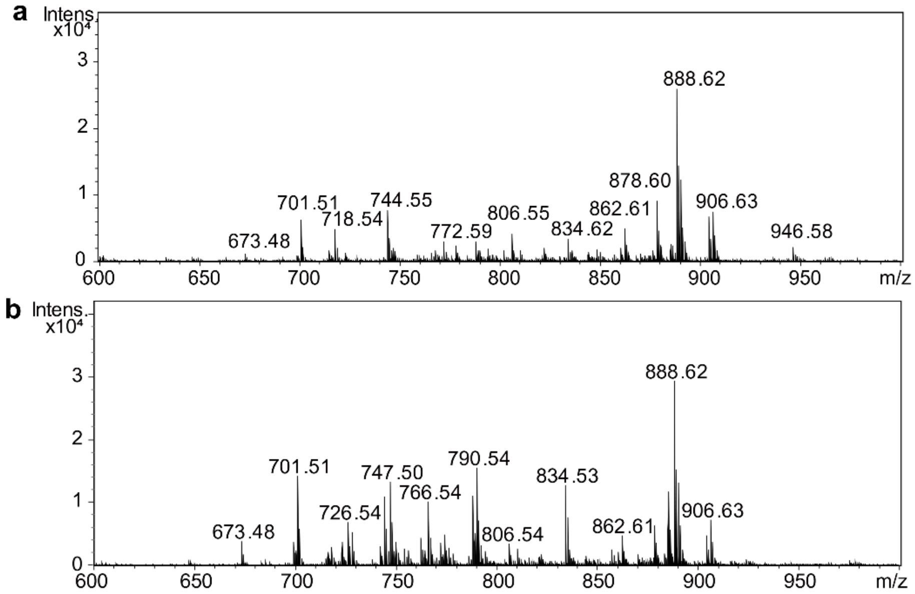


**Supplementary Fig. 4**: Representative mass spectra obtained from a) expanded and b) fresh-frozen mouse cerebrum tissue slices in the negative ion mode using a laser raster spot size of 20 μm.

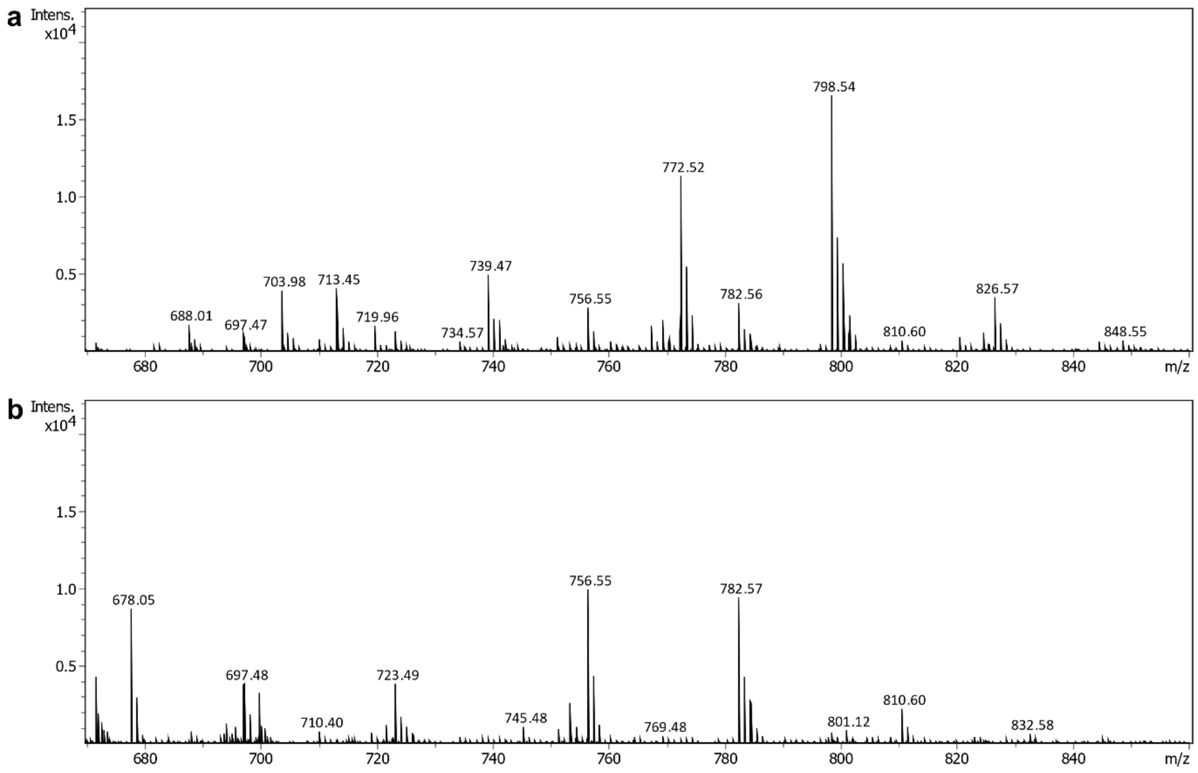


**Supplementary Fig. 5**: Representative mass spectra obtained from a) expanded and b) fresh-frozen mouse cerebrum tissue slices in the positive ion mode using a laser raster spot size of 20 µm.

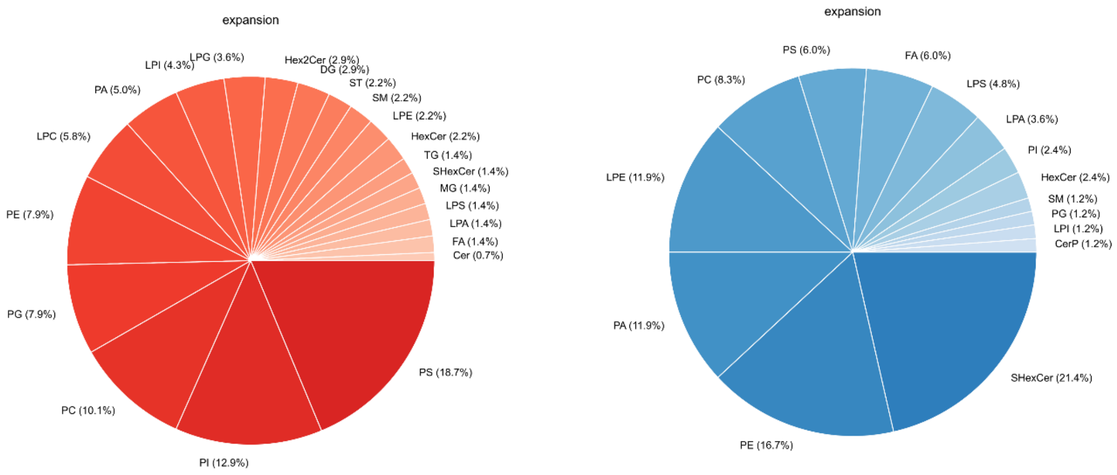


**Supplementary Fig. 6**: Subclass statistical analyses of the identified lipids in expanded mouse brain tissue in positive and negative ion modes.

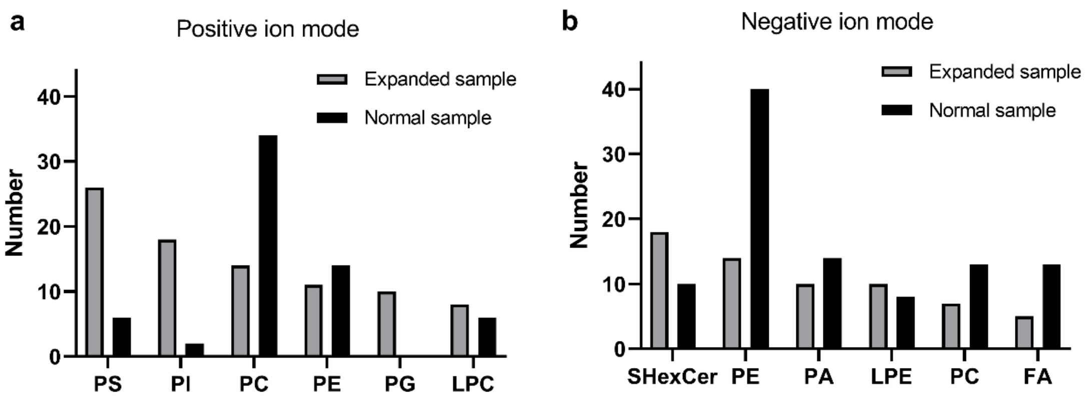


**Supplementary Fig. 7**: Classification of assigned lipid classes (top 6) from expanded mouse cerebrum tissue slices in the a) positive and b) negative ion mode at a single measurement using a laser raster spot size of 10 µm and its comparison with normal samples. Specific assignments are listed in Suppl. Table 1-4.

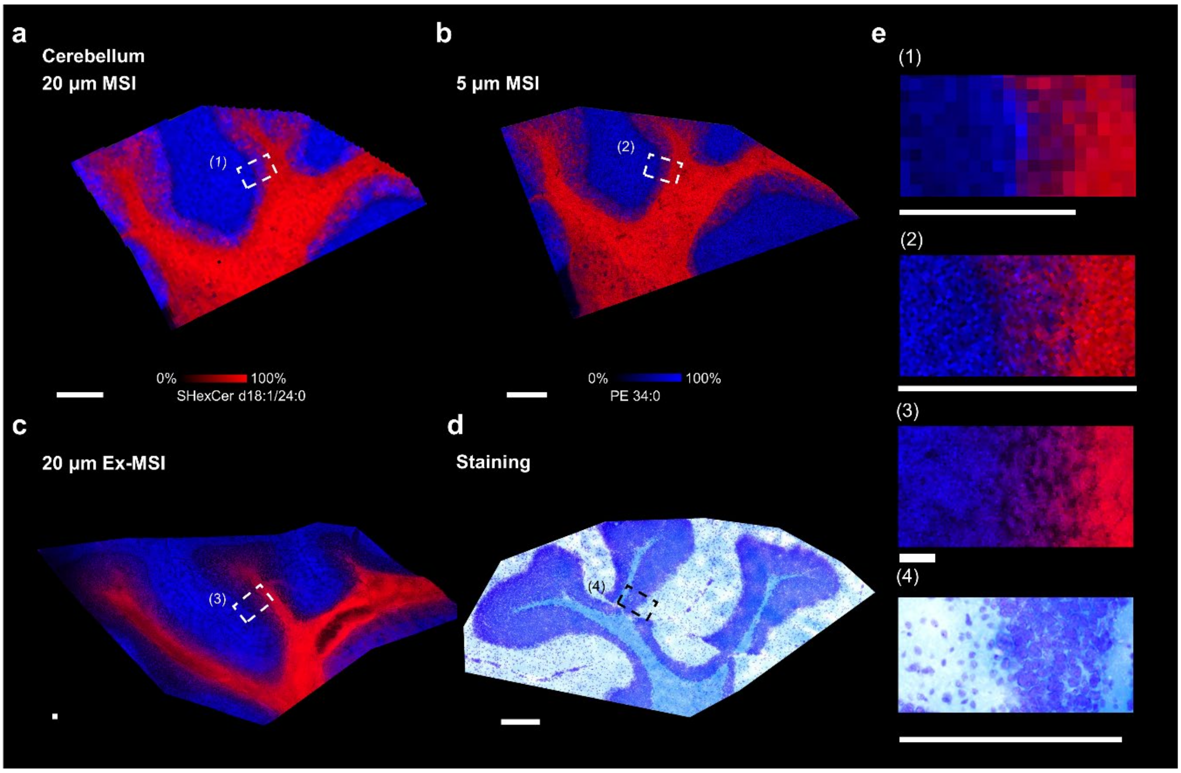


**Supplementary Fig. 8**: overlay ion images of m/z 718.54 (blue) and m/z 888.62 (red) for mouse cerebellum acquired by (a) 20 µm MSI, b) 5 µm MSI, and c) 20 µm 10X Ex-MSI (d). d) f optical image of mouse cerebellum after staining using luxol fast blue and cresyl violet. g) expended view of boxed regiones in (a-d). Scar bar represents 300 µm.

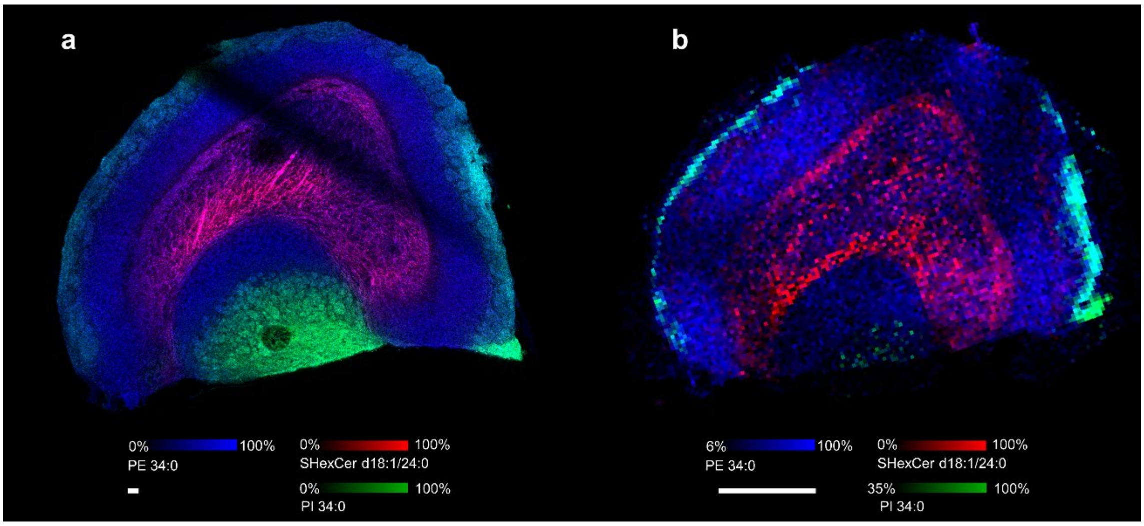


**Supplementary Fig. 9**: overlay ion images of m/z 718.54 (blue) m/z 837.55 (green), and m/z 888.62 (red) for mouse olfactory acquired by a) 20 µm 10X EX-MSI, b) 20 µm MSI. Scar bar represents 500 µm.

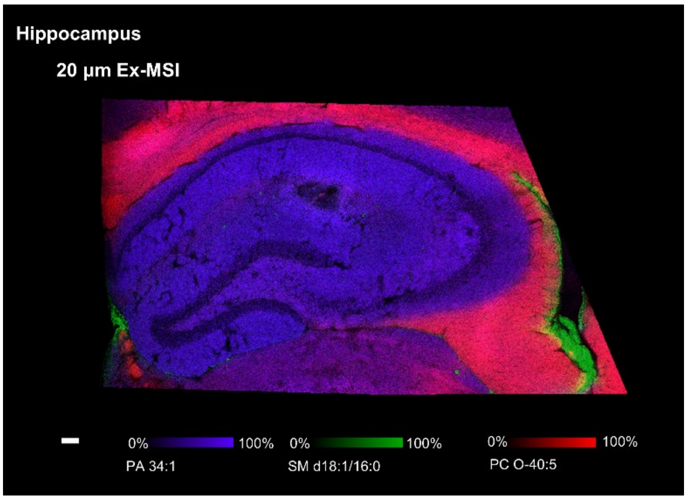


**Supplementary Fig. 10**: overlay ion images of m/z 823.65 (red), m/z 697.48 (blue), and m/z 725.56 (Green) for mouse hippocampus acquired by 20 µm 10X EX-MSI in the positive ion mode.  Scar bar represents 1 mm.

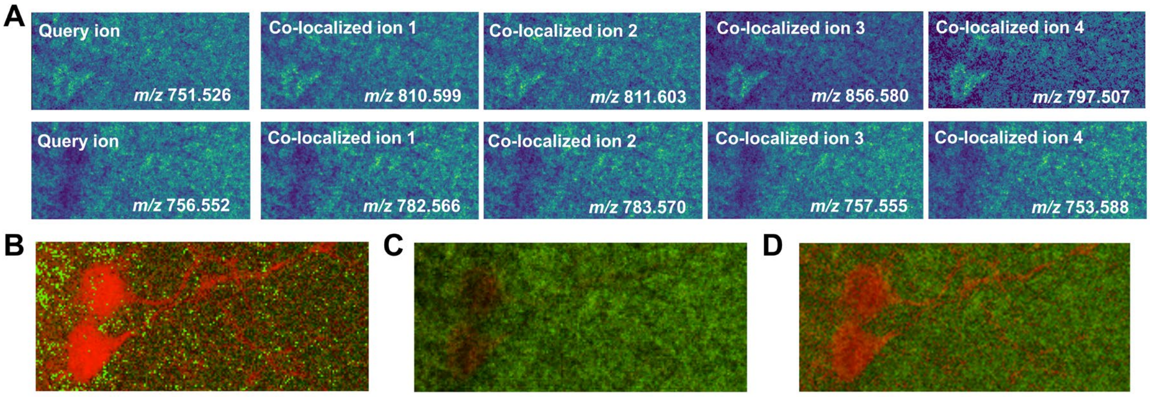


**Supplementary Fig. 11**: Results for enhancing ROI Quality.

**Supplementary Table 1**. List of lipid assignments within 5 ppm mass error for the ion species detected in expanded mouse cerebrum slice under positive ion mode. Laser pixel size was 10 µm.

| Theoretical m/z | Experimental m/z | Error (ppm) | Identity | Chemical Formula | Ion Type |
|---|---|---|---|---|---|
| 373.213 | 373.2114 | 4.3 | FA 22:6 | C22H32O2Na2 | $[M+2Na-H]^+$ |
| 377.2444 | 377.2427 | 4.5 | FA 22:4 | C22H36O2Na2 | $[M+2Na-H]^+$ |
| 441.2385 | 441.2402 | 3.9 | MG 22:6 | C25H38O4K | $[M+K]^+$ |
| 467.3737 | 467.3731 | 1.3 | ST 28:0;O5 | C28H50O5 | $[M+H]^+$ |
| 476.3099 | 476.3111 | 2.5 | LPC O-14:0 | C22H48NO6PNa | $[M+Na]^+$ |
| 518.2622 | 518.2643 | 4.2 | LPE 18:1 | C23H46NO7PK | $[M+K]^+$ |
| 520.2774 | 520.28 | 5.0 | LPE 18:0 | C23H48NO7PK | $[M+K]^+$ |
| 523.2417 | 523.2433 | 3.1 | LPG 16:0 | C22H45O9PK | $[M+K]^+$ |
| 524.3726 | 524.3711 | 2.9 | LPC 18:0 | C26H54NO7P | $[M+H]^+$ |
| 528.3424 | 528.3424 | 0.2 | LPC O-18:2 | C26H52NO6PNa | $[M+Na]^+$ |
| 531.287 | 531.2847 | 4.3 | LPA 22:1 | C25H49O7PK | $[M+K]^+$ |
| 532.2789 | 532.28 | 2.1 | LPC 16:1 | C24H48NO7PK | $[M+K]^+$ |
| 532.374 | 532.3737 | 0.4 | LPC O-18:0 | C26H56NO6PNa | $[M+Na]^+$ |
| 533.3002 | 533.3004 | 0.4 | LPA 22:0 | C25H51O7PK | $[M+K]^+$ |
| 538.2806 | 538.2809 | 0.6 | ST 24:1;O5;T | C26H45NO7SNa | $[M+Na]^+$ |
| 542.25 | 542.2489 | 2.0 | LPS 18:3 | C24H42NO9PNa | $[M+Na]^+$ |
| 556.2795 | 556.28 | 0.9 | LPC 18:3 | C26H48NO7PK | $[M+K]^+$ |
| 564.2677 | 564.2698 | 3.7 | LPS 18:0 | C24H48NO9PK | $[M+K]^+$ |
| 564.307 | 564.3061 | 1.6 | LPC 20:5 | C28H48NO7PNa | $[M+Na]^+$ |
| 573.4854 | 573.4853 | 0.2 | DG O-32:2 | C35H66O4Na | $[M+Na]^+$ |
| 576.3046 | 576.3061 | 2.6 | LPE 24:6 | C29H48NO7PNa | $[M+Na]^+$ |
| 583.3007 | 583.3006 | 0.2 | LPG 22:4 | C28H49O9PNa | $[M+Na]^+$ |
| 605.3206 | 605.3215 | 1.5 | LPG 22:1 | C28H55O9PK | $[M+K]^+$ |
| 609.3153 | 609.3139 | 2.3 | LPG 22:2 | C28H53O9PNa2 | $[M+2Na-H]^+$ |

| | | | | | |
|---|---|---|---|---|---|
| 611.3325 | 611.3295 | 4.7 | LPG 22:1 | C28H55O9PNa2 | $[M+2Na-H]^+$ |
| 612.3018 | 612.3037 | 3.1 | LPC 22:6 | C30H50NO7PNa2 | $[M+2Na-H]^+$ |
| 615.4955 | 615.4959 | 0.6 | DG 34:2 | C37H68O5Na | $[M+Na]^+$ |
| 619.2883 | 619.2854 | 4.7 | LPI 18:2 | C27H49O12PNa | $[M+Na]^+$ |
| 635.3661 | 635.3659 | 0.3 | PA 28:1 | C31H59O8PNa2 | $[M+2Na-H]^+$ |
| 639.2934 | 639.2906 | 4.4 | LPI 18:0 | C27H53O12PK | $[M+K]^+$ |
| 663.2907 | 663.2906 | 0.2 | LPI 20:2 | C29H53O12PK | $[M+K]^+$ |
| 665.2649 | 665.2673 | 3.6 | LPI 20:4 | C29H49O12PNa2 | $[M+2Na-H]^+$ |
| 671.3182 | 671.3167 | 2.2 | LPI 22:4 | C31H53O12PNa | $[M+Na]^+$ |
| 677.3664 | 677.3636 | 4.1 | LPI 22:1 | C31H59O12PNa | $[M+Na]^+$ |
| 681.4838 | 681.4855 | 2.5 | DG 38:5 | C41H70O5K | $[M+K]^+$ |
| 684.4964 | 684.4939 | 3.7 | PC O-28:1 | C36H72NO7PNa | $[M+Na]^+$ |
| 688.4511 | 688.4524 | 1.9 | PS O-28:0 | C34H68NO9PNa | $[M+Na]^+$ |
| 692.5931 | 692.5928 | 0.4 | Cer d18:1/24:1 | C42H81NO3Na2 | $[M+2Na-H]^+$ |
| 697.4775 | 697.4779 | 0.6 | PA 34:1 | C37H71O8PNa | $[M+Na]^+$ |
| 711.4959 | 711.4959 | 0.0 | DG 42:10 | C45H68O5Na | $[M+Na]^+$ |
| 719.463 | 719.4624 | 0.8 | PG O-30:0 | C36H73O9PK | $[M+K]^+$ |
| 723.4938 | 723.4935 | 0.4 | PA 36:2 | C39H73O8PNa | $[M+Na]^+$ |
| 728.5218 | 728.5201 | 2.3 | PC 30:0 | C38H76NO8PNa | $[M+Na]^+$ |
| 730.4435 | 730.442 | 2.1 | PS O-30:1 | C36H70NO9PK | $[M+K]^+$ |
| 732.4571 | 732.4576 | 0.7 | PS O-30:0 | C36H72NO9PK | $[M+K]^+$ |
| 734.5693 | 734.5694 | 0.1 | PC 32:0 | C40H80NO8P | $[M+H]^+$ |
| 738.5065 | 738.5044 | 2.8 | PE 34:2 | C39H74NO8PNa | $[M+Na]^+$ |
| 739.5622 | 739.5612 | 1.4 | PA O-38:1 | C41H81O7PNa | $[M+Na]^+$ |
| 740.5588 | 740.5589 | 0.1 | PC O-34:4 | C42H78NO7P | $[M+H]^+$ |
| 744.517 | 744.515 | 2.7 | PS O-32:0 | C38H76NO9PNa | $[M+Na]^+$ |
| 745.4775 | 745.4779 | 0.5 | PA 38:5 | C41H71O8PNa | $[M+Na]^+$ |
| 748.5838 | 748.5851 | 1.7 | PE 36:0 | C41H82NO8P | $[M+H]^+$ |
| 749.5094 | 749.5092 | 0.3 | PA 38:3 | C41H75O8PNa | $[M+Na]^+$ |
| 758.4989 | 758.4967 | 2.9 | PS 34:3 | C40H72NO10P | $[M+H]^+$ |

| | | | | | |
|---|---|---|---|---|---|
| 760.5243 | 760.5252 | 1.2 | PC O-34:5 | C42H76NO7PNa | $[M+Na]^+$ |
| 761.4611 | 761.4575 | 4.6 | PI O-28:1 | C37H71O12PNa | $[M+Na]^+$ |
| 762.5056 | 762.5044 | 1.6 | PE 36:4 | C41H74NO8PNa | $[M+Na]^+$ |
| 768.5866 | 768.5878 | 1.6 | PC O-34:1 | C42H84NO7PNa | $[M+Na]^+$ |
| 772.526 | 772.5253 | 0.9 | PC 32:0 | C40H80NO8PK | $[M+K]^+$ |
| 772.5615 | 772.5617 | 0.3 | PE O-36:0 | C41H84NO7PK | $[M+K]^+$ |
| 774.5981 | 774.6007 | 3.4 | PE 38:1 | C43H84NO8P | $[M+H]^+$ |
| 778.5733 | 778.5721 | 1.5 | PE O-38:3 | C43H82NO7PNa | $[M+Na]^+$ |
| 780.479 | 780.4786 | 0.5 | PS 34:3 | C40H72NO10PNa | $[M+Na]^+$ |
| 780.5525 | 780.5514 | 1.4 | PC 34:2 | C42H80NO8PNa | $[M+Na]^+$ |
| 781.6206 | 781.6194 | 1.5 | SM d18:1/20:0 | C43H87N2O6PNa | $[M+Na]^+$ |
| 782.5678 | 782.5694 | 2.2 | PC 36:4 | C44H80NO8P | $[M+H]^+$ |
| 783.349 | 782.567 | 0.9 | PC 34:1 | C42H82NO8PNa | $[M+Na]^+$ |
| 783.5029 | 783.5018 | 1.4 | PI 30:0 | C39H75O13P | $[M+H]^+$ |
| 784.5149 | 784.5123 | 3.3 | PS 36:4 | C42H74NO10P | $[M+H]^+$ |
| 787.4861 | 787.4886 | 3.2 | PG 34:1 | C40H77O10PK | $[M+K]^+$ |
| 788.5206 | 788.5202 | 0.5 | PS O-34:0 | C40H80NO9PK | $[M+K]^+$ |
| 789.5777 | 789.5745 | 4.1 | PA O-40:1 | C43H85O7PNa2 | $[M+2Na-H]^+$ |
| 790.5342 | 790.5357 | 1.9 | PE 38:4 | C43H78NO8PNa | $[M+Na]^+$ |
| 794.5479 | 794.546 | 2.4 | PE O-38:3 | C43H82NO7PK | $[M+K]^+$ |
| 802.6305 | 802.632 | 1.9 | PE 40:1 | C45H88NO8P | $[M+H]^+$ |
| 804.5501 | 804.5514 | 1.6 | PC 36:4 | C44H80NO8PNa | $[M+Na]^+$ |
| 804.6297 | 804.6324 | 3.4 | HexCer d18:1/22:1 | C46H87NO8Na | $[M+Na]^+$ |
| 809.5166 | 809.5175 | 1.1 | PI 32:1 | C41H77O13P | $[M+H]^+$ |
| 809.65 | 809.6507 | 0.9 | SM d18:1/22:0 | C45H91N2O6PNa | $[M+Na]^+$ |
| 810.528 | 810.5256 | 3.0 | PS 36:2 | C42H78NO10PNa | $[M+Na]^+$ |
| 812.544 | 812.5412 | 3.3 | PS 36:1 | C42H80NO10PNa | $[M+Na]^+$ |
| 814.5567 | 814.5569 | 0.2 | PS 36:0 | C42H82NO10PNa | $[M+Na]^+$ |
| 816.5761 | 816.5749 | 1.3 | PS 38:2 | C44H82NO10P | $[M+H]^+$ |

| | | | | | |
|---|---|---|---|---|---|
| 820.619 | 820.6191 | 0.1 | PC O-38:3 | C46H88NO7PNa | [M+Na]$^+$ |
| 824.5769 | 824.5776 | 0.8 | PS O-38:2 | C44H84NO9PNa | [M+Na]$^+$ |
| 829.5754 | 829.5743 | 1.3 | TG 48:8 | C51H82O6K | [M+K]$^+$ |
| 832.5125 | 832.5099 | 3.1 | PS 38:5 | C44H76NO10PNa | [M+Na]$^+$ |
| 834.4828 | 834.4798 | 3.6 | SHexCer d18:1/16:0;OH | C40H77NO12SK | [M+K]$^+$ |
| 834.5264 | 834.5256 | 1.0 | PS 38:4 | C44H78NO10PNa | [M+Na]$^+$ |
| 834.634 | 834.6347 | 0.8 | PE O-42:3 | C47H90NO7PNa | [M+Na]$^+$ |
| 835.6686 | 835.6663 | 2.8 | SM d18:1/24:1 | C47H93N2O6PNa | [M+Na]$^+$ |
| 844.5646 | 844.5617 | 3.4 | PE O-42:6 | C47H84NO7PK | [M+K]$^+$ |
| 845.5677 | 845.5668 | 1.1 | PG 38:0 | C44H87O10PK | [M+K]$^+$ |
| 845.6692 | 845.6654 | 4.5 | TG 52:9 | C55H88O6 | [M+H]$^+$ |
| 850.6649 | 850.666 | 1.3 | PC O-40:2 | C48H94NO7PNa | [M+Na]$^+$ |
| 852.4755 | 852.4762 | 0.8 | PS 38:6 | C44H74NO10PNa2 | [M+2Na-H]$^+$ |
| 852.5151 | 852.5151 | 0.0 | PS 38:3 | C44H80NO10PK | [M+K]$^+$ |
| 854.6441 | 854.6456 | 1.8 | HexCer d18:1/24:1 | C48H91NO8Na2 | [M+2Na-H]$^+$ |
| 863.5232 | 863.5199 | 3.8 | PG 40:5 | C46H81O10PK | [M+K]$^+$ |
| 864.5155 | 864.5126 | 3.4 | PS O-40:7 | C46H78NO9PNa2 | [M+2Na-H]$^+$ |
| 864.576 | 864.5725 | 4.0 | PS 40:3 | C46H84NO10PNa | [M+Na]$^+$ |
| 864.64 | 864.6407 | 0.8 | Hex2Cer d18:0/16:0 | C46H89NO13 | [M+H]$^+$ |
| 866.65 | 866.6482 | 2.1 | HexCer d18:1/24:0;OH | C48H93NO9K | [M+K]$^+$ |
| 869.5664 | 869.5668 | 0.6 | PG 40:2 | C46H87O10PK | [M+K]$^+$ |
| 871.5511 | 871.5484 | 3.1 | PG 44:10 | C50H79O10P | [M+H]$^+$ |
| 871.5805 | 871.5825 | 2.3 | PG 40:1 | C46H89O10PK | [M+K]$^+$ |
| 872.6399 | 872.6375 | 2.8 | PS 42:2 | C48H90NO10P | [M+H]$^+$ |
| 878.625 | 878.6245 | 0.6 | PS O-42:3 | C48H90NO9PNa | [M+Na]$^+$ |
| 885.5041 | 885.5042 | 0.1 | PG 42:8 | C48H79O10PK | [M+K]$^+$ |

| | | | | | |
|---|---|---|---|---|---|
| | | | SHexCer | | |
| 890.5467 | 890.5424 | 4.7 | d18:1/20:0;OH | C44H85NO12SK | [M+K]+ |
| 893.5687 | 893.5668 | 2.1 | PG 42:4 | C48H87O10PK | [M+K]+ |
| 895.5815 | 895.5825 | 1.1 | PG 42:3 | C48H89O10PK | [M+K]+ |
| 899.6162 | 899.6138 | 2.7 | PG 42:1 | C48H93O10PK | [M+K]+ |
| 915.5712 | 915.5723 | 1.2 | PI O-38:2 | C47H89O12PK | [M+K]+ |
| 916.6014 | 916.6038 | 2.6 | PS 44:5 | C50H88NO10PNa | [M+Na]+ |
| 920.5609 | 920.5566 | 4.7 | PC 44:10 | C52H84NO8PK | [M+K]+ |
| 922.5019 | 922.4995 | 2.6 | PS 44:10 | C50H78NO10PK | [M+K]+ |
| 925.5176 | 925.5203 | 2.9 | PI 38:4 | C47H83O13PK | [M+K]+ |
| 927.5362 | 927.5359 | 0.3 | PI 38:3 | C47H85O13PK | [M+K]+ |
| 932.576 | 932.5777 | 1.8 | PS 44:5 | C50H88NO10PK | [M+K]+ |
| 934.5949 | 934.5934 | 1.6 | PS 44:4 | C50H90NO10PK | [M+K]+ |
| 936.6064 | 936.609 | 2.8 | PS 44:3 | C50H92NO10PK | [M+K]+ |
| 939.5328 | 939.5334 | 0.6 | PI O-40:7 | C49H83O12PNa2 | [M+2Na-H]]+ |
| | | | Hex2Cer | | |
| 940.6649 | 940.6696 | 5.0 | d18:1/20:0 | C50H95NO13Na | [M+Na]+ |
| 945.5784 | 945.5827 | 4.5 | PI O-42:7 | C51H87O12PNa | [M+Na]+ |
| 949.5192 | 949.5203 | 1.2 | PI 40:6 | C49H83O13PK | [M+K]+ |
| 955.5697 | 955.5672 | 2.6 | PI 40:3 | C49H89O13PK | [M+K]+ |
| 967.5655 | 967.5647 | 0.8 | PI O-42:7 | C51H87O12PNa2 | [M+2Na-H]+ |
| 973.5198 | 973.5203 | 0.5 | PI 42:8 | C51H83O13PK | [M+K]+ |
| 975.5321 | 975.5359 | 3.9 | PI 42:7 | C51H85O13PK | [M+K]+ |
| 975.6931 | 975.6896 | 3.6 | PI 44:2 | C53H99O13P | [M+H]+ |
| 981.5796 | 981.5829 | 3.4 | PI 42:4 | C51H91O13PK | [M+K]+ |
| 997.5243 | 997.5203 | 4.0 | PI 44:10 | C53H83O13PK | [M+K]+ |
| 1001.549 | 1001.552 | 2.6 | PI 44:8 | C53H87O13PK | [M+K]+ |
| 1007.597 | 1007.599 | 1.3 | PI 44:5 | C53H93O13PK | [M+K]+ |
| | | | Hex2Cer | | |
| 1024.759 | 1024.764 | 4.1 | d18:1/26:0 | C56H107NO13Na | [M+Na]+ |

| | | | | | |
|---|---|---|---|---|---|
| | | | Hex2Cer | | |
| 1046.742 | 1046.745 | 3.0 | d18:1/26:0 | C56H107NO13Na2 | [M+2Na-H]+ |

**Supplementary Table 2**. List of lipid assignments within 5 ppm mass error for the ion species detected in expanded mouse cerebrum slice under negative ion mode. Laser pixel size was 10 µm.

| Theoretical m/z | Experimental m/z | Error (ppm) | Identity | Chemical Formula | Ion Type |
|---|---|---|---|---|---|
| 255.2326 | 255.233 | 1.6 | FA 16:0 | C16H32O2 | $[M-H]^-$ |
| 281.2485 | 281.2486 | 0.4 | FA 18:1 | C18H34O2 | $[M-H]^-$ |
| 283.2639 | 283.2643 | 1.4 | FA 18:0 | C18H36O2 | $[M-H]^-$ |
| 303.2327 | 303.233 | 1.0 | FA 20:4 | C20H32O2 | $[M-H]^-$ |
| 327.2329 | 327.233 | 0.3 | FA 22:6 | C22H32O2 | $[M-H]^-$ |
| 409.2362 | 409.2361 | 0.2 | LPA 16:0 | C19H39O7P | $[M-H]^-$ |
| 435.2513 | 435.2517 | 0.9 | LPA 18:1 | C21H41O7P | $[M-H]^-$ |
| 437.2665 | 437.2674 | 2.1 | LPA 18:0 | C21H43O7P | $[M-H]^-$ |
| 452.2791 | 452.2783 | 1.8 | LPE 16:0 | C21H44NO7P | $[M-H]^-$ |
| 460.2849 | 460.2834 | 3.3 | LPE O-18:3 | C23H44NO6P | $[M-H]^-$ |
| 462.2974 | 462.299 | 3.5 | LPE O-18:2 | C23H46NO6P | $[M-H]^-$ |
| 478.2933 | 478.2939 | 1.3 | LPE 18:1 | C23H46NO7P | $[M-H]^-$ |
| 480.3084 | 480.3096 | 2.5 | LPE 18:0 | C23H48NO7P | $[M-H]^-$ |
| 490.329 | 490.3303 | 2.7 | LPE O-20:2 | C25H50NO6P | $[M-H]^-$ |
| 506.2894 | 506.2888 | 1.2 | LPS O-18:2 | C24H46NO8P | $[M-H]^-$ |
| 506.3259 | 506.3252 | 1.4 | LPE 20:1 | C25H50NO7P | $[M-H]^-$ |
| 508.3417 | 508.3409 | 1.6 | LPE 20:0 | C25H52NO7P | $[M-H]^-$ |
| 524.3002 | 524.2994 | 1.5 | LPS 18:0 | C24H48NO9P | $[M-H]^-$ |
| 538.2702 | 538.2706 | 0.7 | LPE 20:3 | C25H46NO7P | $[M+Cl]^-$ |
| 540.2841 | 540.2863 | 4.1 | LPE 20:2 | C25H48NO7P | $[M+Cl]^-$ |
| 566.2653 | 566.2655 | 0.4 | LPS O-20:4 | C26H46NO8P | $[M+Cl]^-$ |
| 582.2578 | 582.2604 | 4.5 | LPS 20:3 | C26H46NO9P | $[M+Cl]^-$ |
| 599.3192 | 599.3202 | 1.7 | LPI 18:0 | C27H53O12P | $[M-H]^-$ |
| 644.5013 | 644.5025 | 1.9 | CerP d18:1/18:0 | C36H72NO6P | $[M-H]^-$ |

| | | | | | |
|---|---|---|---|---|---|
| 647.4637 | 647.4657 | 3.1 | PA 32:0 | C35H69O8P | [M-H]- |
| 673.4814 | 673.4814 | 0.0 | PA 34:1 | C37H71O8P | [M-H]- |
| 675.4987 | 675.497 | 2.5 | PA 34:0 | C37H73O8P | [M-H]- |
| 699.4955 | 699.497 | 2.1 | PA 36:2 | C39H73O8P | [M-H]- |
| 701.5115 | 701.5127 | 1.7 | PA 36:1 | C39H75O8P | [M-H]- |
| 716.5219 | 716.5236 | 2.4 | PE 34:1 | C39H76NO8P | [M-H]- |
| 718.5387 | 718.5392 | 0.7 | PE 34:0 | C39H78NO8P | [M-H]- |
| 729.5428 | 729.544 | 1.6 | PA 38:1 | C41H79O8P | [M-H]- |
| 731.442 | 731.4424 | 0.5 | PA 36:4 | C39H69O8P | [M+Cl]- |
| 732.441 | 732.4377 | 4.5 | PC 30:4 | C38H68NO8P | [M+Cl]- |
| 744.5552 | 744.5549 | 0.4 | PE 36:1 | C41H80NO8P | [M-H]- |
| 747.4969 | 747.497 | 0.1 | PA 40:6 | C43H73O8P | [M-H]- |
| 748.467 | 748.469 | 2.7 | PE 34:3 | C39H72NO8P | [M+Cl]- |
| 757.4551 | 757.4581 | 4.0 | PA 38:5 | C41H71O8P | [M+Cl]- |
| 759.4724 | 759.4737 | 1.7 | PA 38:4 | C41H73O8P | [M+Cl]- |
| 762.4838 | 762.4846 | 1.0 | PC 32:3 | C40H74NO8P | [M+Cl]- |
| 762.5658 | 762.5656 | 0.3 | HexCer d18:1/18:0 | C42H81NO8 | [M+Cl]- |
| 764.464 | 764.4624 | 2.1 | SHexCer d18:1/14:1;OH | C38H71NO12S | [M-H]- |
| 765.5676 | 765.5683 | 0.9 | SM d18:1/18:0 | C41H83N2O6P | [M+Cl]- |
| 766.4774 | 766.4781 | 0.9 | SHexCer d18:1/14:0;OH | C38H73NO12S | [M-H]- |
| 766.5378 | 766.5392 | 1.8 | PE 38:4 | C43H78NO8P | [M-H]- |
| 772.5845 | 772.5862 | 2.2 | PE 38:1 | C43H84NO8P | [M-H]- |
| 776.4989 | 776.5003 | 1.8 | PE 36:3 | C41H76NO8P | [M+Cl]- |
| 778.4766 | 778.4795 | 3.7 | PS O-34:3 | C40H74NO9P | [M+Cl]- |
| 784.4437 | 784.4442 | 0.6 | SHexCer d18:1/14:1;OH | C38H71NO11S | [M+Cl]- |
| 788.5427 | 788.5447 | 2.5 | PS 36:1 | C42H80NO10P | [M-H]- |
| 790.5411 | 790.5392 | 2.4 | PE 40:6 | C45H78NO8P | [M-H]- |
| 792.4951 | 792.4937 | 1.8 | SHexCer d18:1/16:1;OH | C40H75NO12S | [M-H]- |
| 794.4718 | 794.4744 | 3.3 | PS 34:2 | C40H74NO10P | [M+Cl]- |
| 802.5147 | 802.5159 | 1.5 | PE 38:4 | C43H78NO8P | [M+Cl]- |

| | | | | | |
|---|---|---|---|---|---|
| 806.5471 | 806.5472 | 0.1 | PE 38:2 | C43H82NO8P | [M+Cl]$^-$ |
| 806.5901 | 806.5836 | 8.1 | PC O-36:2 | C44H86NO7P | [M+Cl]$^-$ |
| 820.5242 | 820.525 | 1.0 | SHexCer d18:1/18:1;OH | C42H79NO12S | [M-H]$^-$ |
| 820.6065 | 820.5993 | 8.9 | PE O-40:2 | C45H88NO7P | [M+Cl]$^-$ |
| 822.5376 | 822.5407 | 3.8 | SHexCer d18:1/18:0;OH | C42H81NO12S | [M-H]$^-$ |
| 830.5471 | 830.5472 | 0.1 | PE 40:4 | C45H82NO8P | [M+Cl]$^-$ |
| 832.6022 | 832.5993 | 3.6 | PC O-38:3 | C46H88NO7P | [M+Cl]$^-$ |
| 834.5742 | 834.5771 | 3.5 | SHexCer d18:1/20:0 | C44H85NO11S | [M-H]$^-$ |
| 835.5335 | 835.5342 | 0.8 | PI 34:1 | C43H81O13P | [M-H]$^-$ |
| 837.5452 | 837.5418 | 4.1 | PG 38:2 | C44H83O10P | [M+Cl]$^-$ |
| 846.5034 | 846.5057 | 2.7 | PS 38:4 | C44H78NO10P | [M+Cl]$^-$ |
| 848.6376 | 848.6306 | 8.4 | PE O-42:2 | C47H92NO7P | [M+Cl]$^-$ |
| 850.6357 | 850.6331 | 3.1 | PE 44:4 | C49H90NO8P | [M-H]$^-$ |
| 850.5708 | 850.572 | 1.4 | SHexCer d18:1/20:0; OH | C44H85NO12S | [M-H]$^-$ |
| 860.5893 | 860.5927 | 4.0 | SHexCer d18:1/22:1 | C46H87NO11S | [M-H]$^-$ |
| 860.6377 | 860.6388 | 1.3 | HexCer 42:2;O3 | C48H91NO9 | [M+Cl]$^-$ |
| 862.6076 | 862.6084 | 0.9 | SHexCer d18:1/22:0 | C46H89NO11S | [M-H]$^-$ |
| 874.6068 | 874.6098 | 3.4 | PC 40:3 | C48H90NO8P | [M+Cl]$^-$ |
| 876.5845 | 876.5876 | 3.5 | SHexCer d18:1/22:1;OH | C46H87NO12S | [M-H]$^-$ |
| 876.6226 | 876.6255 | 3.3 | PC 40:2 | C48H92NO8P | [M+Cl]$^-$ |
| 878.6037 | 878.6033 | 0.5 | SHexCer d18:1/22:0;OH | C46H89NO12S | [M-H]$^-$ |
| 885.5491 | 885.5499 | 0.9 | PI 38:4 | C47H83O13P | [M-H]$^-$ |
| 886.6066 | 886.6084 | 2.0 | SHexCer d18:1/24:1 | C48H89NO11S | [M-H]$^-$ |
| 888.6226 | 888.624 | 1.6 | SHexCer d18:1/24:0 | C48H91NO11S | [M-H]$^-$ |
| 890.6356 | 890.6397 | 4.6 | SHexCer d18:0/24:0 | C48H93NO11S | [M-H]$^-$ |
| 892.6197 | 892.6204 | 0.8 | PS O-42:2 | C48H92NO9P | [M+Cl]$^-$ |
| 902.6006 | 902.6033 | 3.0 | SHexCer d18:1/24:2;OH | C48H89NO12S | [M-H]$^-$ |
| 902.6387 | 902.6411 | 2.7 | PC 42:3 | C50H94NO8P | [M+Cl]$^-$ |
| 904.616 | 904.6189 | 3.2 | SHexCer d18:1/24:1;OH | C48H91NO12S | [M-H]$^-$ |

| | | | | | |
|---|---|---|---|---|---|
| 906.6328 | 906.6346 | 2.0 | SHexCer d18:1/24:0;OH | C48H93NO12S | $[M-H]^-$ |

**Supplementary Table 3**. List of lipid assignments within 5 ppm mass error for the ion species detected in the fresh-frozen mouse cerebrum slice under positive ion mode. Laser pixel size was 10 μm.

| Theoretical m/z | Experimental m/z | Error (ppm) | Identity | Chemical Formula | Ion Type |
|---|---|---|---|---|---|
| 301.212 | 301.2114 | 2 | FA 16:0 | C16H32O2Na2 | $[M+2Na-H]^+$ |
| 327.2268 | 327.227 | 0.6 | FA 18:1 | C18H34O2Na2 | $[M+2Na-H]^+$ |
| 329.2427 | 329.2427 | 0 | FA 18:0 | C18H36O2Na2 | $[M+2Na-H]^+$ |
| 349.2111 | 349.2114 | 0.9 | FA 20:4 | C20H32O2Na2 | $[M+2Na-H]^+$ |
| 367.1211 | 367.121 | 0.3 | ST 18:4;O3;S | C18H22O6S | $[M+H]^+$ |
| 373.2113 | 373.2114 | 0.3 | FA 22:6 | C22H32O2Na2 | $[M+2Na-H]^+$ |
| 415.2219 | 415.222 | 0.2 | CPA 16:0 | C19H37O6PNa | $[M+Na]^+$ |
| 419.2577 | 419.2557 | 4.8 | CPA 18:1 | C21H39O6P | $[M+H]^+$ |
| 431.1938 | 431.1959 | 4.9 | CPA 16:0 | C19H37O6PK | $[M+K]^+$ |
| 437.2036 | 437.2039 | 0.7 | CPA 16:0 | C19H37O6PNa2 | $[M+2Na-H]^+$ |
| 441.2372 | 441.2376 | 0.9 | CPA 18:1 | C21H39O6PNa | $[M+Na]^+$ |
| 443.2522 | 443.2533 | 2.5 | CPA 18:0 | C21H41O6PNa | $[M+Na]^+$ |
| 455.2134 | 455.2145 | 2.4 | LPA 16:0 | C19H39O7PNa2 | $[M+2Na-H]^+$ |
| 457.2325 | 457.2326 | 0.2 | LPA 18:2 | C21H39O7PNa | $[M+Na]^+$ |
| 459.2467 | 459.2482 | 3.3 | LPA 18:1 | C21H41O7PNa | $[M+Na]^+$ |
| 463.218 | 463.2196 | 3.5 | CPA 18:1 | C21H39O6PNa2 | $[M+2Na-H]^+$ |
| 465.2335 | 465.2352 | 3.7 | CPA 18:0 | C21H41O6PNa2 | $[M+2Na-H]^+$ |
| 469.2673 | 469.2665 | 1.7 | LPA O-18:0 | C21H45O6PNa2 | $[M+2Na-H]^+$ |
| 481.2304 | 481.2326 | 4.6 | LPA 20:4 | C23H39O7PNa | $[M+Na]^+$ |
| 487.2782 | 487.2795 | 2.7 | LPA 20:1 | C23H45O7PNa | $[M+Na]^+$ |
| 492.2478 | 492.2487 | 1.8 | LPE 16:0 | C21H44NO7PK | $[M+K]^+$ |

| | | | | | |
|---|---|---|---|---|---|
| 496.3384 | 496.3398 | 2.8 | LPC 16:0 | C24H50NO7P | [M+H]⁺ |
| 504.3439 | 504.3424 | 3 | LPC O-16:0 | C24H52NO6PNa | [M+Na]⁺ |
| 505.228 | 505.2302 | 4.4 | LPA 20:3 | C23H41O7PNa2 | [M+2Na-H]⁺ |
| 505.243 | 505.2408 | 4.4 | ST 19:1;O3;GlcA | C25H38O9Na | [M+Na]⁺ |
| 509.2603 | 509.2615 | 2.4 | LPA 20:1 | C23H45O7PNa2 | [M+2Na-H]⁺ |
| 516.308 | 516.3061 | 3.7 | LPC 16:1 | C24H48NO7PNa | [M+Na]⁺ |
| 518.3218 | 518.3217 | 0.2 | LPC 16:0 | C24H50NO7PNa | [M+Na]⁺ |
| 524.37 | 524.3711 | 2.1 | LPC 18:0 | C26H54NO7P | [M+H]⁺ |
| 526.2926 | 526.2928 | 0.4 | LPE 22:6 | C27H44NO7P | [M+H]⁺ |
| 546.3505 | 546.353 | 4.6 | LPC 18:0 | C26H54NO7PNa | [M+Na]⁺ |
| 560.333 | 560.3323 | 1.2 | LPS O-20:1 | C26H52NO8PNa | [M+Na]⁺ |
| 568.2801 | 568.28 | 0.2 | LPE 22:4 | C27H48NO7PK | [M+K]⁺ |
| 575.5002 | 575.501 | 1.4 | FAHFA 35:0;O | C35H68O4Na | [M+Na]⁺ |
| 583.2955 | 583.2982 | 4.6 | LPG 20:1 | C26H51O9PNa2 | [M+2Na-H]⁺ |
| 585.3125 | 585.3139 | 2.4 | LPG 20:0 | C26H53O9PNa2 | [M+2Na-H]⁺ |
| 602.3429 | 602.3428 | 0.2 | LPS 22:1 | C28H54NO9PNa | [M+Na]⁺ |
| 613.3448 | 613.3452 | 0.7 | LPG 22:0 | C28H57O9PNa2 | [M+2Na-H]⁺ |
| 655.4849 | 655.4785 | 9.8 | CerPE 32:1;O2 | C34H69N2O6PNa | [M+Na]⁺ |
| 673.4247 | 673.4229 | 2.7 | DG 38:9 | C41H62O5K | [M+K]⁺ |
| 689.4149 | 689.4129 | 2.9 | PA 32:2 | C35H65O8PNa2 | [M+2Na-H]⁺ |
| 693.4446 | 693.4442 | 0.6 | PA 32:0 | C35H69O8PNa2 | [M+2Na-H]⁺ |
| 695.4614 | 695.4622 | 1.2 | PA 34:2 | C37H69O8PNa | [M+Na]⁺ |
| 696.4593 | 696.4575 | 2.6 | PC 28:2 | C36H68NO8PNa | [M+Na]⁺ |
| 697.4779 | 697.4779 | 0 | PA 34:1 | C37H71O8PNa | [M+Na]⁺ |
| 700.4887 | 700.4888 | 0.1 | PC 28:0 | C36H72NO8PNa | [M+Na]⁺ |
| 701.455 | 701.4542 | 1.1 | DG 40:9 | C43H66O5K | [M+K]⁺ |
| 711.4832 | 711.4838 | 0.8 | SM 32:2;O2 | C37H73N2O6PK | [M+K]⁺ |
| 712.4647 | 712.4678 | 4.4 | PE O-32:2 | C37H72NO7PK | [M+K]⁺ |
| 713.4498 | 713.4518 | 2.8 | PA 34:1 | C37H71O8PK | [M+K]⁺ |
| 715.4271 | 715.4285 | 2 | PA 34:3 | C37H67O8PNa2 | [M+2Na-H]⁺ |

| | | | | | |
|---|---|---|---|---|---|
| 716.4996 | 716.4991 | 0.7 | PE O-32:0 | C37H76NO7PK | [M+K]⁺ |
| 717.449 | 717.4466 | 3.3 | PA 36:5 | C39H67O8PNa | [M+Na]⁺ |
| 719.4597 | 719.4598 | 0.1 | PA 34:1 | C37H71O8PNa2 | [M+2Na-H]⁺ |
| 721.4748 | 721.4755 | 1 | PA 34:0 | C37H73O8PNa2 | [M+2Na-H]⁺ |
| 723.4939 | 723.4935 | 0.6 | PA 36:2 | C39H73O8PNa | [M+Na]⁺ |
| 723.5401 | 723.5411 | 1.4 | SM 34:2;O2 | C39H77N2O6PNa | [M+Na]⁺ |
| 725.5554 | 725.5568 | 1.9 | SM 34:1;O2 | C39H79N2O6PNa | [M+Na]⁺ |
| 728.5209 | 728.5225 | 2.2 | PC 32:3 | C40H74NO8P | [M+H]⁺ |
| 733.4739 | 733.478 | 5.6 | Mannosyl-1beta-phosphomycoketide C31 | C37H75O9PK | [M+K]⁺ |
| 733.5749 | 733.5742 | 1 | PA 38:0 | C41H81O8P | [M+H]⁺ |
| 734.5697 | 734.5694 | 0.4 | PC 32:0 | C40H80NO8P | [M+H]⁺ |
| 739.4296 | 739.4285 | 1.5 | PA 36:5 | C39H67O8PNa2 | [M+2Na-H]⁺ |
| 741.4416 | 741.4442 | 3.5 | PA 36:4 | C39H69O8PNa2 | [M+2Na-H]⁺ |
| 742.5714 | 742.5721 | 0.9 | PC O-32:0 | C40H82NO7PNa | [M+Na]⁺ |
| 743.4614 | 743.4598 | 2.2 | PA 36:3 | C39H71O8PNa2 | [M+2Na-H]⁺ |
| 745.4768 | 745.4755 | 1.7 | PA 36:2 | C39H73O8PNa2 | [M+2Na-H]⁺ |
| 747.4878 | 747.4911 | 4.4 | PA 36:1 | C39H75O8PNa2 | [M+2Na-H]⁺ |
| 748.4831 | 748.4864 | 4.4 | PC 30:1 | C38H74NO8PNa2 | [M+2Na-H]⁺ |
| 749.5062 | 749.5068 | 0.8 | PA 36:0 | C39H77O8PNa2 | [M+2Na-H]⁺ |
| 750.5873 | 750.5854 | 2.5 | HexCer 36:1;O2 | C42H81NO8Na | [M+Na]⁺ |
| 751.525 | 751.5248 | 0.3 | PA 38:2 | C41H77O8PNa | [M+Na]⁺ |
| 751.5739 | 751.5724 | 2 | SM 36:2;O2 | C41H81N2O6PNa | [M+Na]⁺ |
| 753.5893 | 753.5881 | 1.6 | SM 36:1;O2 | C41H83N2O6PNa | [M+Na]⁺ |
| 754.5356 | 754.5357 | 0.1 | PC 32:1 | C40H78NO8PNa | [M+Na]⁺ |
| 755.537 | 755.5351 | 2.5 | PA O-38:1 | C41H81O7PK | [M+K]⁺ |
| 756.5512 | 756.5514 | 0.3 | PC 32:0 | C40H80NO8PNa | [M+Na]⁺ |
| 757.5542 | 757.5508 | 4.5 | PA O-38:0 | C41H83O7PK | [M+K]⁺ |
| 760.5857 | 760.5851 | 0.8 | PC 34:1 | C42H82NO8P | [M+H]⁺ |

| | | | | | |
|---|---|---|---|---|---|
| 762.5013 | 762.502 | 0.9 | PE 34:1 | C39H76NO8PNa2 | $[M+2Na-H]^+$ |
| 764.5181 | 764.5177 | 0.5 | PE 34:0 | C39H78NO8PNa2 | $[M+2Na-H]^+$ |
| 768.591 | 768.5902 | 1 | PC O-36:4 | C44H82NO7P | $[M+H]^+$ |
| 769.474 | 769.4755 | 1.9 | PA 38:4 | C41H73O8PNa2 | $[M+2Na-H]^+$ |
| 770.6069 | 770.6058 | 1.4 | PC O-36:3 | C44H84NO7P | $[M+H]^+$ |
| 771.4945 | 771.4935 | 1.3 | PA 40:6 | C43H73O8PNa | $[M+Na]^+$ |
| 772.5217 | 772.5253 | 4.7 | PC 32:0 | C40H80NO8PK | $[M+K]^+$ |
| 773.5114 | 773.5092 | 2.8 | PA 40:5 | C43H75O8PNa | $[M+Na]^+$ |
| 776.5145 | 776.5177 | 4.1 | PC 32:1 | C40H78NO8PNa2 | $[M+2Na-H]^+$ |
| 780.4913 | 780.494 | 3.5 | PE 36:3 | C41H76NO8PK | $[M+K]^+$ |
| 781.6221 | 781.6194 | 3.5 | SM 38:1;O2 | C43H87N2O6PNa | $[M+Na]^+$ |
| 782.5665 | 782.567 | 0.6 | PC 34:1 | C42H82NO8PNa | $[M+Na]^+$ |
| 784.5807 | 784.5827 | 2.5 | PC 34:0 | C42H84NO8PNa | $[M+Na]^+$ |
| 784.6175 | 784.6191 | 2 | PE O-38:0 | C43H88NO7PNa | $[M+Na]^+$ |
| 786.6 | 786.6007 | 0.9 | PC 36:2 | C44H84NO8P | $[M+H]^+$ |
| 788.6158 | 788.6164 | 0.8 | PC 36:1 | C44H86NO8P | $[M+H]^+$ |
| 790.5346 | 790.5357 | 1.4 | PE 38:4 | C43H78NO8PNa | $[M+Na]^+$ |
| 793.4763 | 793.4755 | 1 | PA 40:6 | C43H73O8PNa2 | $[M+2Na-H]^+$ |
| 795.4905 | 795.4911 | 0.8 | PA 40:5 | C43H75O8PNa2 | $[M+2Na-H]^+$ |
| 796.581 | 796.5827 | 2.1 | PE 38:1 | C43H84NO8PNa | $[M+Na]^+$ |
| 797.51 | 797.5068 | 4 | PA 40:4 | C43H77O8PNa2 | $[M+2Na-H]^+$ |
| 799.5653 | 799.5614 | 4.9 | PA 40:0 | C43H85O8PK | $[M+K]^+$ |
| 804.5514 | 804.5514 | 0 | PC 36:4 | C44H80NO8PNa | $[M+Na]^+$ |
| 808.4837 | 808.4864 | 3.3 | PE 38:6 | C43H74NO8PNa2 | $[M+2Na-H]^+$ |
| 808.5039 | 808.5075 | 4.5 | PS 34:0 | C40H78NO10PNa2 | $[M+2Na-H]^+$ |
| 808.5805 | 808.5827 | 2.7 | PC 36:2 | C44H84NO8PNa | $[M+Na]^+$ |
| 809.6508 | 809.6507 | 0.1 | SM 40:1;O2 | C45H91N2O6PNa | $[M+Na]^+$ |
| 810.5979 | 810.5983 | 0.5 | PC 36:1 | C44H86NO8PNa | $[M+Na]^+$ |
| 812.5148 | 812.5177 | 3.6 | PE 38:4 | C43H78NO8PNa2 | $[M+2Na-H]^+$ |
| 813.5965 | 813.598 | 1.8 | TG 46:5 | C49H84O6Na2 | $[M+2Na-H]^+$ |

| | | | | | |
|---|---|---|---|---|---|
| 818.5609 | 818.5646 | 4.5 | PE 38:1 | C43H84NO8PNa2 | [M+2Na-H]$^+$ |
| 820.5217 | 820.5253 | 4.4 | PC 36:4 | C44H80NO8PK | [M+K]$^+$ |
| 822.5369 | 822.541 | 5 | PC 36:3 | C44H82NO8PK | [M+K]$^+$ |
| 824.5166 | 824.5177 | 1.3 | PC 36:5 | C44H78NO8PNa2 | [M+2Na-H]$^+$ |
| 826.5722 | 826.5723 | 0.1 | PC 36:1 | C44H86NO8PK | [M+K]$^+$ |
| 828.5488 | 828.549 | 0.2 | PC 36:3 | C44H82NO8PNa2 | [M+2Na-H]$^+$ |
| 830.5629 | 830.5646 | 2 | PC 36:2 | C44H84NO8PNa2 | [M+2Na-H]$^+$ |
| 832.5818 | 832.5827 | 1.1 | PC 38:4 | C46H84NO8PNa | [M+Na]$^+$ |
| 836.5166 | 836.5177 | 1.3 | PE 40:6 | C45H78NO8PNa2 | [M+2Na-H]$^+$ |
| 836.6512 | 836.6504 | 1 | PE O-42:2 | C47H92NO7PNa | [M+Na]$^+$ |
| 840.5137 | 840.5126 | 1.3 | PS O-38:5 | C44H78NO9PNa2 | [M+2Na-H]$^+$ |
| 840.5504 | 840.5514 | 1.2 | PE 42:7 | C47H80NO8PNa | [M+Na]$^+$ |
| 848.52 | 848.5201 | 0.1 | PC 40:10 | C48H76NO8PNa | [M+Na]$^+$ |
| 848.5359 | 848.5388 | 3.4 | PT 36:1 | C43H82NO10PNa2 | [M+2Na-H]$^+$ |
| 850.5289 | 850.5263 | 3.1 | Hex2Cer 30:1;O2 | C42H79NO13Na2 | [M+2Na-H]$^+$ |
| 852.5492 | 852.549 | 0.2 | PC 38:5 | C46H82NO8PNa2 | [M+2Na-H]$^+$ |
| 856.578 | 856.5803 | 2.7 | PC 38:3 | C46H86NO8PNa2 | [M+2Na-H]$^+$ |
| 858.5257 | 858.5256 | 0.1 | PS 40:6 | C46H78NO10PNa | [M+Na]$^+$ |
| 858.5931 | 858.5959 | 3.3 | PC 38:2 | C46H88NO8PNa2 | [M+2Na-H]$^+$ |
| 868.5282 | 868.5253 | 3.3 | PC 40:8 | C48H80NO8PK | [M+K]$^+$ |
| 878.5564 | 878.5576 | 1.4 | Hex2Cer 32:1;O2 | C44H83NO13Na2 | [M+2Na-H]$^+$ |
| 880.5081 | 880.5075 | 0.7 | PS 40:6 | C46H78NO10PNa2 | [M+2Na-H]$^+$ |
| 880.5789 | 880.5803 | 1.6 | PC 40:5 | C48H86NO8PNa2 | [M+2Na-H]$^+$ |
| 902.488 | 902.4918 | 4.2 | PS 42:9 | C48H76NO10PNa2 | [M+2Na-H]$^+$ |
| 903.4954 | 903.497 | 1.8 | PI 36:4 | C45H79O13PNa2 | [M+2Na-H]$^+$ |
| 908.5768 | 908.5777 | 1 | PS 42:3 | C48H88NO10PK | [M+K]$^+$ |
| 912.6396 | 912.6429 | 3.6 | PC 42:3 | C50H94NO8PNa2 | [M+2Na-H]$^+$ |
| 952.6139 | 952.613 | 0.9 | SHexCer 42:1;O3 | C48H93NO12SNa2 | [M+2Na-H]$^+$ |
| 993.6359 | 993.6378 | 1.9 | PI 42:1 | C51H97O13PNa2 | [M+2Na-H]$^+$ |

**Supplementary Table 4**. List of lipid assignments within 5 ppm mass error for the ion species detected in the fresh-frozen mouse cerebrum slice under negative ion mode. Laser pixel size was 10 µm.

| Theoretical m/z | Experimental m/z | Error (ppm) | Identity | Chemical Formula | Ion Type |
|---|---|---|---|---|---|
| 255.2331 | 255.233 | 0.391799 | FA 16:0 | C16H32O2 | [M-H]- |
| 279.2329 | 279.233 | 0.358124 | FA 18:2 | C18H32O2 | [M-H]- |
| 281.2487 | 281.2486 | 0.355557 | FA 18:1 | C18H34O2 | [M-H]- |
| 283.2643 | 283.2643 | 0 | FA 18:0 | C18H36O2 | [M-H]- |
| 303.233 | 303.233 | 0 | FA 20:4 | C20H32O2 | [M-H]- |
| 305.2482 | 305.2486 | 1.310407 | FA 20:3 | C20H34O2 | [M-H]- |
| 307.2642 | 307.2643 | 0.325453 | FA 20:2 | C20H36O2 | [M-H]- |
| 309.2802 | 309.2799 | 0.969995 | FA 20:1 | C20H38O2 | [M-H]- |
| 311.2956 | 311.2956 | 0 | FA 20:0 | C20H40O2 | [M-H]- |
| 321.1249 | 321.1263 | 4.359655 | ST 18:4;O3 | C18H22O3 | [M+Cl]- |
| 327.2336 | 327.233 | 1.833556 | FA 22:6 | C22H32O2 | [M-H]- |
| 331.2646 | 331.2643 | 0.905621 | FA 22:4 | C22H36O2 | [M-H]- |
| 337.3118 | 337.3112 | 1.778773 | FA 22:1 | C22H42O2 | [M-H]- |
| 339.3262 | 339.3269 | 2.062907 | FA 22:0 | C22H44O2 | [M-H]- |
| 409.2355 | 409.2361 | 1.466146 | LPA 16:0 | C19H39O7P | [M-H]- |
| 417.241 | 417.2412 | 0.479339 | CPA 18:1 | C21H39O6P | [M-H]- |
| 419.2562 | 419.2568 | 1.431104 | CPA 18:0 | C21H41O6P | [M-H]- |
| 427.2013 | 427.2022 | 2.106731 | CPA 16:0 | C19H37O6P | [M+Cl]- |
| 435.2523 | 435.2517 | 1.378513 | LPA 18:1 | C21H41O7P | [M-H]- |
| 436.2845 | 436.2834 | 2.521297 | LPE O-16:1 | C21H44NO6P | [M-H]- |
| 437.2672 | 437.2674 | 0.457386 | LPA 18:0 | C21H43O7P | [M-H]- |
| 463.2821 | 463.283 | 1.942657 | LPA 20:1 | C23H45O7P | [M-H]- |
| 478.2938 | 478.2939 | 0.209076 | LPE 18:1 | C23H46NO7P | [M-H]- |
| 483.2749 | 483.2729 | 4.138448 | LPG 16:0 | C22H45O9P | [M-H]- |
| 500.2787 | 500.2783 | 0.80 | LPE 20:4 | C25H44NO7P | [M-H]- |

| | | | | | |
|---|---|---|---|---|---|
| 506.3259 | 506.3252 | 1.38 | LPE 20:1 | C25H50NO7P | [M-H]- |
| 507.2713 | 507.2729 | -3.15 | LPG 18:2 | C24H45O9P | [M-H]- |
| 508.3404 | 508.3409 | -0.98 | LPE 20:0 | C25H52NO7P | [M-H]- |
| 509.2871 | 509.2885 | -2.75 | LPG 18:1 | C24H47O9P | [M-H]- |
| 524.2773 | 524.2783 | -1.91 | LPE 22:6 | C27H44NO7P | [M-H]- |
| 524.3004 | 524.2994 | 1.91 | LPS 18:0 | C24H48NO9P | [M-H]- |
| 528.3093 | 528.3096 | -0.57 | LPE 22:4 | C27H48NO7P | [M-H]- |
| 540.2867 | 540.2863 | 0.74 | LPE 20:2 | C25H48NO7P | [M+Cl]- |
| 547.267 | 547.2678 | -1.46 | PA 23:3;O2 | C26H45O10P | [M-H]- |
| 574.3165 | 574.3151 | 2.44 | LPC 20:4;O2 | C28H50NO9P | [M-H]- |
| 581.3115 | 581.3096 | 3.27 | PG 21:1;O | C27H51O11P | [M-H]- |
| 599.3197 | 599.3202 | -0.83 | LPI 18:0 | C27H53O12P | [M-H]- |
| 673.4822 | 673.4814 | 1.19 | PA 34:1 | C37H71O8P | [M-H]- |
| 687.5008 | 687.4994 | 2.04 | DG 42:10 | C45H68O5 | [M-H]- |
| 688.495 | 688.4923 | 3.92 | PE 32:1 | C37H72NO8P | [M-H]- |
| 692.4048 | 692.4064 | -2.31 | PE 30:3 | C35H64NO8P | [M+Cl]- |
| 699.4965 | 699.497 | -0.71 | PA 36:2 | C39H73O8P | [M-H]- |
| 700.5296 | 700.5287 | 1.28 | PE O-34:2 | C39H76NO7P | [M-H]- |
| 701.5125 | 701.5127 | -0.29 | PA 36:1 | C39H75O8P | [M-H]- |
| 702.4383 | 702.4352 | 4.41 | PS 30:2 | C36H66NO10P | [M-H]- |
| 715.5751 | 715.576 | -1.26 | SM 35:1;O2 | C40H81N2O6P | [M-H]- |
| 716.5247 | 716.5236 | 1.54 | PE 34:1 | C39H76NO8P | [M-H]- |
| 718.5408 | 718.5392 | 2.23 | PE 34:0 | C39H78NO8P | [M-H]- |
| 719.4649 | 719.4657 | -1.11 | PA 38:6 | C41H69O8P | [M-H]- |
| 722.4899 | 722.4897 | 0.28 | PC O-30:2 | C38H74NO7P | [M+Cl]- |
| 723.498 | 723.497 | 1.38 | PA 38:4 | C41H73O8P | [M-H]- |
| 725.5112 | 725.5127 | -2.07 | PA 38:3 | C41H75O8P | [M-H]- |
| 726.5455 | 726.5443 | 1.65 | PE O-36:3 | C41H78NO7P | [M-H]- |
| 728.5603 | 728.56 | 0.41 | PE O-36:2 | C41H80NO7P | [M-H]- |
| 737.4625 | 737.461 | 2.03 | PI O-28:1 | C37H71O12P | [M-H]- |

| | | | | | |
|---|---|---|---|---|---|
| 738.5076 | 738.5079 | -0.41 | PE 36:4 | C41H74NO8P | [M-H]- |
| 742.5427 | 742.5392 | 4.71 | PE 36:2 | C41H78NO8P | [M-H]- |
| 744.554 | 744.5549 | -1.21 | PE 36:1 | C41H80NO8P | [M-H]- |
| 745.4801 | 745.4814 | -1.74 | PA 40:7 | C43H71O8P | [M-H]- |
| 746.5097 | 746.513 | -4.42 | PE O-38:7 | C43H74NO7P | [M-H]- |
| 747.4982 | 747.497 | 1.61 | PA 40:6 | C43H73O8P | [M-H]- |
| 748.5278 | 748.5287 | -1.20 | PE O-38:6 | C43H76NO7P | [M-H]- |
| 750.5433 | 750.5443 | -1.33 | PE O-38:5 | C43H78NO7P | [M-H]- |
| 753.5592 | 753.5571 | 2.79 | PA O-38:0 | C41H83O7P | [M+Cl]- |
| 754.5772 | 754.5756 | 2.12 | PE O-38:3 | C43H82NO7P | [M-H]- |
| 755.5599 | 755.5596 | 0.40 | PA 40:2 | C43H81O8P | [M-H]- |
| 756.5917 | 756.5913 | 0.53 | PE O-38:2 | C43H84NO7P | [M-H]- |
| 759.4746 | 759.4737 | 1.19 | PA 38:4 | C41H73O8P | [M+Cl]- |
| 762.4826 | 762.4846 | -2.62 | PC 32:3 | C40H74NO8P | [M+Cl]- |
| 762.5101 | 762.5079 | 2.89 | PE 38:6 | C43H74NO8P | [M-H]- |
| 762.5644 | 762.5656 | -1.57 | HexCer 36:1;O2 | C42H81NO8 | [M+Cl]- |
| 764.522 | 764.5236 | -2.09 | PE 38:5 | C43H76NO8P | [M-H]- |
| 765.5686 | 765.5683 | 0.39 | SM 36:1;O2 | C41H83N2O6P | [M+Cl]- |
| 766.5386 | 766.5392 | -0.78 | PE 38:4 | C43H78NO8P | [M-H]- |
| 770.5689 | 770.5705 | -2.08 | PE 38:2 | C43H82NO8P | [M-H]- |
| 772.5288 | 772.5287 | 0.13 | PE O-40:8 | C45H76NO7P | [M-H]- |
| 772.5868 | 772.5862 | 0.78 | PE 38:1 | C43H84NO8P | [M-H]- |
| 773.5362 | 773.5338 | 3.10 | PG 36:2 | C42H79O10P | [M-H]- |
| 774.545 | 774.5443 | 0.90 | PE O-40:7 | C45H78NO7P | [M-H]- |
| 775.4137 | 775.4111 | 3.35 | PA 40:10 | C43H65O8P | [M+Cl]- |
| 775.4429 | 775.4403 | 3.35 | PI 30:3 | C39H69O13P | [M-H]- |
| 776.5601 | 776.56 | 0.13 | PE O-40:6 | C45H80NO7P | [M-H]- |
| 778.5783 | 778.5756 | 3.47 | PE O-40:5 | C45H82NO7P | [M-H]- |
| 780.5711 | 780.568 | 3.97 | PC O-34:1 | C42H84NO7P | [M+Cl]- |
| 780.5924 | 780.5913 | 1.41 | PE O-40:4 | C45H84NO7P | [M-H]- |

| | | | | | |
|---|---|---|---|---|---|
| 783.5874 | 783.5909 | -4.47 | PA 42:2 | C45H85O8P | [M-H]- |
| 786.5274 | 786.5291 | -2.16 | PS 36:2 | C42H78NO10P | [M-H]- |
| 788.5443 | 788.5447 | -0.51 | PS 36:1 | C42H80NO10P | [M-H]- |
| 790.54 | 790.5392 | 1.01 | PE 40:6 | C45H78NO8P | [M-H]- |
| 794.5493 | 794.5472 | 2.64 | PC 34:1 | C42H82NO8P | [M+Cl]- |
| 794.5697 | 794.5705 | -1.01 | PE 40:4 | C45H82NO8P | [M-H]- |
| 795.4896 | 795.4948 | -6.54 | PG 35:2 | C41H77O10P | [M+Cl]- |
| 795.5174 | 795.5182 | -1.01 | PG 38:5 | C44H77O10P | [M-H]- |
| 795.5276 | 795.5312 | -4.53 | PG O-36:2 | C42H81O9P | [M+Cl]- |
| 796.5502 | 796.5498 | 0.50 | PS O-38:4 | C44H80NO9P | [M-H]- |
| 798.5985 | 798.6018 | -4.13 | PE 40:2 | C45H86NO8P | [M-H]- |
| 799.658 | 799.6588 | -1.00 | TG 45:0 | C48H92O6 | [M+Cl]- |
| 800.6159 | 800.6175 | -2.00 | PE 40:1 | C45H88NO8P | [M-H]- |
| 802.5159 | 802.5159 | 0.00 | PE 38:4 | C43H78NO8P | [M+Cl]- |
| 804.5294 | 804.5316 | -2.73 | PE 38:3 | C43H80NO8P | [M+Cl]- |
| 806.5483 | 806.5458 | 3.10 | SHexCer 36:1;O2 | C42H81NO11S | [M-H]- |
| 807.5062 | 807.5029 | 4.09 | PI 32:1 | C41H77O13P | [M-H]- |
| 808.5162 | 808.5134 | 3.46 | PS 38:5 | C44H76NO10P | [M-H]- |
| 809.5103 | 809.5105 | -0.25 | PG 36:2 | C42H79O10P | [M+Cl]- |
| 810.5293 | 810.5291 | 0.25 | PS 38:4 | C44H78NO10P | [M-H]- |
| 814.5597 | 814.5604 | -0.86 | PS 38:2 | C44H82NO10P | [M-H]- |
| 816.5748 | 816.576 | -1.47 | PS 38:1 | C44H84NO10P | [M-H]- |
| 820.5592 | 820.5629 | -4.51 | PC 36:2 | C44H84NO8P | [M+Cl]- |
| 820.6001 | 820.5993 | 0.97 | PE O-40:2 | C45H88NO7P | [M+Cl]- |
| 821.5433 | 821.5469 | -4.38 | PG O-38:3 | C44H83O9P | [M+Cl]- |
| 822.5436 | 822.5407 | 3.53 | SHexCer 36:1;O3 | C42H81NO12S | [M-H]- |
| 822.5791 | 822.5785 | 0.73 | PC 36:1 | C44H86NO8P | [M+Cl]- |
| 824.5805 | 824.5811 | -0.73 | PS O-40:4 | C46H84NO9P | [M-H]- |
| 824.5972 | 824.5942 | 3.64 | PC 36:0 | C44H88NO8P | [M+Cl]- |
| 834.5295 | 834.5291 | 0.48 | PS 40:6 | C46H78NO10P | [M-H]- |

| | | | | | |
|---|---|---|---|---|---|
| 834.5769 | 834.5771 | -0.24 | SHexCer 38:1;O2 | C44H85NO11S | [M-H]- |
| 834.5769 | 834.5785 | -1.92 | PE 40:2 | C45H86NO8P | [M+Cl]- |
| 836.6183 | 836.6175 | 0.96 | PC 40:4 | C48H88NO8P | [M-H]- |
| 836.6344 | 836.6306 | 4.54 | PC O-38:1 | C46H92NO7P | [M+Cl]- |
| 837.5409 | 837.5418 | -1.07 | PG 38:2 | C44H83O10P | [M+Cl]- |
| 839.5593 | 839.5574 | 2.26 | PG 38:1 | C44H85O10P | [M+Cl]- |
| 841.394 | 841.3912 | 3.33 | PI 30:4;O2 | C39H67O15P | [M+Cl]- |
| 842.5744 | 842.5705 | 4.63 | PE 44:8 | C49H82NO8P | [M-H]- |
| 842.5943 | 842.5917 | 3.09 | PS 40:2 | C46H86NO10P | [M-H]- |
| 844.5985 | 844.5993 | -0.95 | PE O-42:4 | C47H88NO7P | [M+Cl]- |
| 844.643 | 844.6439 | -1.07 | HexCer 42:2;O2 | C48H91NO8 | [M+Cl]- |
| 849.5758 | 849.5782 | -2.82 | PG O-40:3 | C46H87O9P | [M+Cl]- |
| 850.5705 | 850.572 | -1.76 | SHexCer 38:1;O3 | C44H85NO12S | [M-H]- |
| 857.5188 | 857.5186 | 0.23 | PI 36:4 | C45H79O13P | [M-H]- |
| 858.6119 | 858.6149 | -3.49 | PC O-40:4 | C48H90NO7P | [M+Cl]- |
| 860.5912 | 860.5942 | -3.49 | PE 42:3 | C47H88NO8P | [M+Cl]- |
| 861.5457 | 861.5499 | -4.87 | PI 36:2 | C45H83O13P | [M-H]- |
| 862.5763 | 862.5734 | 3.36 | PS O-40:3 | C46H86NO9P | [M+Cl]- |
| 862.6099 | 862.6084 | 1.74 | SHexCer 40:1;O2 | C46H89NO11S | [M-H]- |
| 862.6099 | 862.6098 | 0.12 | PE 42:2 | C47H90NO8P | [M+Cl]- |
| 863.565 | 863.5655 | -0.58 | PI 36:1 | C45H85O13P | [M-H]- |
| 864.623 | 864.6255 | -2.89 | PE 42:1 | C47H92NO8P | [M+Cl]- |
| 865.5802 | 865.5812 | -1.16 | PI 36:0 | C45H87O13P | [M-H]- |
| 874.6083 | 874.6098 | -1.72 | PC 40:3 | C48H90NO8P | [M+Cl]- |
| 876.6242 | 876.6255 | -1.48 | PC 40:2 | C48H92NO8P | [M+Cl]- |
| 878.6043 | 878.6033 | 1.14 | SHexCer 40:1;O3 | C46H89NO12S | [M-H]- |
| 883.5365 | 883.5342 | 2.60 | PI 38:5 | C47H81O13P | [M-H]- |
| 885.5495 | 885.5499 | -0.45 | PI 38:4 | C47H83O13P | [M-H]- |
| 886.6077 | 886.6098 | -2.37 | PE 44:4 | C49H90NO8P | [M+Cl]- |
| 888.6246 | 888.624 | 0.68 | SHexCer 42:2;O2 | C48H91NO11S | [M-H]- |

| | | | | | |
|---|---|---|---|---|---|
| 890.6376 | 890.6397 | -2.36 | SHexCer 42:1;O2 | C48H93NO11S | $[M-H]^-$ |
| 892.6244 | 892.6204 | 4.48 | PS O-42:2 | C48H92NO9P | $[M+Cl]^-$ |
| 902.6416 | 902.6411 | 0.55 | PC 42:3 | C50H94NO8P | $[M+Cl]^-$ |
| 904.6185 | 904.6189 | -0.44 | SHexCer 42:2;O3 | C48H91NO12S | $[M-H]^-$ |
| 906.6326 | 906.6346 | -2.21 | SHexCer 42:1;O3 | C48H93NO12S | $[M-H]^-$ |

**Supplementary Table 5**. List of lipid assignments within 5 ppm mass error for the ion species detected in the A549 cell lines under positive ion mode. Laser pixel size was 5 µm.

| Experimental m/z | Theoretical m/z | Error (ppm) | Identity | Chemical Formula | Ion Type |
|---|---|---|---|---|---|
| 494.3228 | 494.3241 | 2.6 | LPC 16:1 | C24H48NO7P | $[M+H]^+$ |
| 496.3413 | 496.3398 | 3.0 | LPC 16:0 | C24H50NO7P | $[M+H]^+$ |
| 695.4652 | 695.4622 | 4.3 | PA 34:2 | C37H69O8PNa | $[M+Na]^+$ |
| 701.5596 | 701.5592 | 0.6 | SM d18:1/16:1 | C39H77N2O6P | $[M+H]^+$ |
| 703.576 | 703.5748 | 1.7 | SM d18:1/16:0 | C39H79N2O6P | $[M+H]^+$ |
| 704.5212 | 704.5225 | 1.8 | PC 30:1 | C38H74NO8P | $[M+H]^+$ |
| 705.5876 | 705.5905 | 4.1 | SM d18:0/16:0 | C39H81N2O6P | $[M+H]^+$ |
| 706.5372 | 706.5381 | 1.3 | PC 30:0 | C38H76NO8P | $[M+H]^+$ |
| 718.5361 | 718.5381 | 2.8 | PE 34:1 | C39H76NO8P | $[M+H]^+$ |
| 718.5738 | 718.5745 | 1.0 | PC O-32:1 | C40H80NO7P | $[M+H]^+$ |
| 720.5526 | 720.5538 | 1.7 | PE 34:0 | C39H78NO8P | $[M+H]^+$ |
| 720.5868 | 720.5902 | 4.7 | PC O-32:0 | C40H82NO7P | $[M+H]^+$ |
| 721.4773 | 721.4779 | 0.8 | PA 36:3 | C39H71O8PNa | $[M+Na]^+$ |
| 723.493 | 723.4935 | 0.7 | PA 36:2 | C39H73O8PNa | $[M+Na]^+$ |
| 725.5561 | 725.5568 | 1.0 | SM d18:1/16:0 | C39H79N2O6PNa | $[M+Na]^+$ |
| 726.5054 | 726.5068 | 1.9 | PC 32:4 | C40H72NO8P | $[M+H]^+$ |
| 729.5212 | 729.5195 | 2.3 | PA O-36:0 | C39H79O7PK | $[M+K]^+$ |
| 730.5377 | 730.5381 | 0.5 | PC 32:2 | C40H76NO8P | $[M+H]^+$ |
| 731.6051 | 731.6061 | 1.4 | SM d18:1/18:0 | C41H83N2O6P | $[M+H]^+$ |
| 732.5531 | 732.5538 | 1.0 | PC 32:1 | C40H78NO8P | $[M+H]^+$ |
| 734.4529 | 734.4521 | 1.1 | PE O-34:5 | C39H70NO7PK | $[M+K]^+$ |
| 734.5686 | 734.5694 | 1.1 | PC 32:0 | C40H80NO8P | $[M+H]^+$ |
| 740.5557 | 740.5565 | 1.1 | PC O-32:1 | C40H80NO7PNa | $[M+Na]^+$ |
| 744.5879 | 744.5902 | 3.1 | PC O-34:2 | C42H82NO7P | $[M+H]^+$ |
| 746.53 | 746.533 | 4.0 | PS O-34:2 | C40H76NO9P | $[M+H]^+$ |
| 746.5674 | 746.5694 | 2.7 | PE 36:1 | C41H80NO8P | $[M+H]^+$ |

| | | | | | |
|---|---|---|---|---|---|
| 746.6047 | 746.6058 | 1.5 | PC O-34:1 | C42H84NO7P | $[M+H]^+$ |
| 748.5484 | 748.5487 | 0.4 | PS O-34:1 | C40H78NO9P | $[M+H]^+$ |
| 748.5821 | 748.5851 | 4.0 | PE 36:0 | C41H82NO8P | $[M+H]^+$ |
| 750.5628 | 750.5643 | 2.0 | PS O-34:0 | C40H80NO9P | $[M+H]^+$ |
| 754.5363 | 754.5357 | 0.8 | PC 32:1 | C40H78NO8PNa | $[M+Na]^+$ |
| 758.5687 | 758.5694 | 0.9 | PC 34:2 | C42H80NO8P | $[M+H]^+$ |
| 760.5835 | 760.5851 | 2.1 | PC 34:1 | C42H82NO8P | $[M+H]^+$ |
| 762.4816 | 762.4834 | 2.4 | PE O-36:5 | C41H74NO7PK | $[M+K]^+$ |
| 768.5867 | 768.5902 | 4.6 | PC O-36:4 | C44H82NO7P | $[M+H]^+$ |
| 770.5298 | 770.5306 | 1.0 | PS O-34:1 | C40H78NO9PNa | $[M+Na]^+$ |
| 772.5454 | 772.5463 | 1.2 | PS O-34:0 | C40H80NO9PNa | $[M+Na]^+$ |
| 772.5841 | 772.5851 | 1.3 | PE 38:2 | C43H82NO8P | $[M+H]^+$ |
| 774.5625 | 774.5643 | 2.3 | PS O-36:2 | C42H80NO9P | $[M+H]^+$ |
| 774.5993 | 774.6007 | 1.8 | PE 38:1 | C43H84NO8P | $[M+H]^+$ |
| 776.4955 | 776.4991 | 4.6 | PC O-34:5 | C42H76NO7PK | $[M+K]^+$ |
| 776.579 | 776.58 | 1.3 | PS O-36:1 | C42H82NO9P | $[M+H]^+$ |
| 780.5516 | 780.5538 | 2.8 | PC 36:5 | C44H78NO8P | $[M+H]^+$ |
| 782.566 | 782.5694 | 4.3 | PC 36:4 | C44H80NO8P | $[M+H]^+$ |
| 784.5818 | 784.5851 | 4.2 | PC 36:3 | C44H82NO8P | $[M+H]^+$ |
| 786.5988 | 786.6007 | 2.4 | PC 36:2 | C44H84NO8P | $[M+H]^+$ |
| 787.6673 | 787.6687 | 1.8 | SM d18:1/22:0 | C45H91N2O6P | $[M+H]^+$ |
| 788.6131 | 788.6164 | 4.2 | PC 36:1 | C44H86NO8P | $[M+H]^+$ |
| 790.514 | 790.5147 | 0.9 | PE O-38:5 | C43H78NO7PK | $[M+K]^+$ |
| 792.5204 | 792.5174 | 3.8 | PS O-38:7 | C44H74NO9P | $[M+H]^+$ |
| 792.5739 | 792.5749 | 1.3 | PS 36:0 | C42H82NO10P | $[M+H]^+$ |
| 798.5606 | 798.5619 | 1.6 | PS O-36:1 | C42H82NO9PNa | $[M+Na]^+$ |
| 802.5934 | 802.5956 | 2.7 | PS O-38:2 | C44H84NO9P | $[M+H]^+$ |
| 804.5501 | 804.5514 | 1.6 | PC 36:4 | C44H80NO8PNa | $[M+Na]^+$ |
| 806.564 | 806.5624 | 2.0 | Hex2Cer d18:1/12:0 | C42H79NO13 | $[M+H]^+$ |

| | | | | | |
|---|---|---|---|---|---|
| 808.583 | 808.5851 | 2.6 | PC 38:5 | C46H82NO8P | [M+H]+ |
| 810.5968 | 810.6007 | 4.8 | PC 38:4 | C46H84NO8P | [M+H]+ |
| 813.6829 | 813.6844 | 1.8 | SM d18:1/24:1 | C47H93N2O6P | [M+H]+ |
| 814.6294 | 814.632 | 3.2 | PC 38:2 | C46H88NO8P | [M+H]+ |
| 815.6973 | 815.7 | 3.3 | SM d18:1/24:0 | C47H95N2O6P | [M+H]+ |
| 816.5276 | 816.5304 | 3.4 | PE O-40:6 | C45H80NO7PK | [M+K]+ |
| 818.5449 | 818.546 | 1.3 | PE O-40:5 | C45H82NO7PK | [M+K]+ |
| 820.5487 | 820.5463 | 2.9 | PS O-38:4 | C44H80NO9PNa | [M+Na]+ |
| 832.5812 | 832.5851 | 4.7 | PC 40:7 | C48H82NO8P | [M+H]+ |
| 834.5964 | 834.5937 | 3.2 | Hex2Cer d18:1/14:0 | C44H83NO13 | [M+H]+ |
| 835.6654 | 835.6663 | 1.1 | SM d18:1/24:1 | C47H93N2O6PNa | [M+Na]+ |
| 837.6797 | 837.682 | 2.7 | SM d18:1/24:0 | C47H95N2O6PNa | [M+Na]+ |

**Supplementary Table 6**. List of lipid assignments within 5 ppm mass error for the ion species detected in the A549 cell lines under negative ion mode. Laser pixel size was 5 μm.

| Experimental m/z | Theoretical m/z | Error (ppm) | Identity | Chemical Formula | Ion Type |
|---|---|---|---|---|---|
| 253.2183 | 253.2173 | 3.9 | FA 16:1 | C16H30O2 | $[M-H]^-$ |
| 279.2338 | 279.233 | 2.9 | FA 18:2 | C18H32O2 | $[M-H]^-$ |
| 295.2276 | 295.2279 | 1.0 | FA 18:2;O | C18H32O3 | $[M-H]^-$ |
| 297.2443 | 297.2435 | 2.7 | FA 18:1;O | C18H34O3 | $[M-H]^-$ |
| 301.2165 | 301.2173 | 2.7 | FA 20:5 | C20H30O2 | $[M-H]^-$ |
| 303.2331 | 303.233 | 0.3 | FA 20:4 | C20H32O2 | $[M-H]^-$ |
| 305.2494 | 305.2486 | 2.6 | FA 20:3 | C20H34O2 | $[M-H]^-$ |
| 327.2327 | 327.233 | 0.9 | FA 22:6 | C22H32O2 | $[M-H]^-$ |
| 616.4708 | 616.4712 | 0.6 | CerP 34:1;O2 | C34H68NO6P | $[M-H]^-$ |
| 645.4489 | 645.4501 | 1.9 | PA 32:1 | C35H67O8P | $[M-H]^-$ |
| 647.4626 | 647.4657 | 4.8 | PA 32:0 | C35H69O8P | $[M-H]^-$ |
| 671.4644 | 671.4657 | 1.9 | PA 34:2 | C37H69O8P | $[M-H]^-$ |
| 673.4818 | 673.4814 | 0.6 | PA 34:1 | C37H71O8P | $[M-H]^-$ |
| 699.4948 | 699.497 | 3.1 | PA 36:2 | C39H73O8P | $[M-H]^-$ |
| 701.5105 | 701.5127 | 3.1 | PA 36:1 | C39H75O8P | $[M-H]^-$ |
| 716.5216 | 716.5236 | 2.8 | PE 34:1 | C39H76NO8P | $[M-H]^-$ |
| 718.5361 | 718.5392 | 4.3 | PE 34:0 | C39H78NO8P | $[M-H]^-$ |
| 742.5393 | 742.5392 | 0.1 | PE 36:2 | C41H78NO8P | $[M-H]^-$ |
| 744.5544 | 744.5549 | 0.7 | PE 36:1 | C41H80NO8P | $[M-H]^-$ |
| 770.5703 | 770.5705 | 0.3 | PE 38:2 | C43H82NO8P | $[M-H]^-$ |
| 772.5841 | 772.5862 | 2.7 | PE 38:1 | C43H84NO8P | $[M-H]^-$ |
| 885.5471 | 885.5499 | 3.2 | PI 38:4 | C47H83O13P | $[M-H]^-$ |
| 887.5618 | 887.5655 | 4.2 | PI 38:3 | C47H85O13P | $[M-H]^-$ |
